# Archaic Introgression Shaped Human Circadian Traits

**DOI:** 10.1101/2023.02.03.527061

**Authors:** Keila Velazquez-Arcelay, Laura L. Colbran, Evonne McArthur, Colin Brand, David Rinker, Justin Siemann, Douglas McMahon, John A. Capra

**Affiliations:** Department of Biological Sciences, Vanderbilt University; Department of Genetics, Perelman School of Medicine, University of Pennsylvania; Vanderbilt University School of Medicine; Department of Epidemiology and Biostatistics, University of California, San Francisco; Bakar Computational Health Sciences Institute, University of California, San Francisco

**Keywords:** circadian biology, chronotype, Neanderthals, adaptive introgression, gene expression, adaptive evolution

## Abstract

**Introduction:** When the ancestors of modern Eurasians migrated out of Africa and interbred with Eurasian archaic hominins, namely Neanderthals and Denisovans, DNA of archaic ancestry integrated into the genomes of anatomically modern humans. This process potentially accelerated adaptation to Eurasian environmental factors, including reduced ultra-violet radiation and increased variation in seasonal dynamics. However, whether these groups differed substantially in circadian biology, and whether archaic introgression adaptively contributed to human chronotypes remains unknown.

**Results:** Here we traced the evolution of chronotype based on genomes from archaic hominins and present-day humans. First, we inferred differences in circadian gene sequences, splicing, and regulation between archaic hominins and modern humans. We identified 28 circadian genes containing variants with potential to alter splicing in archaics (e.g., *CLOCK*, *PER2*, *RORB*, *RORC*), and 16 circadian genes likely divergently regulated between present-day humans and archaic hominins, including *RORA*. These differences suggest the potential for introgression to modify circadian gene expression. Testing this hypothesis, we found that introgressed variants are enriched among eQTLs for circadian genes. Supporting the functional relevance of these regulatory effects, we found that many introgressed alleles have associations with chronotype. Strikingly, the strongest introgressed effects on chronotype increase morningness, consistent with adaptations to high latitude in other species. Finally, we identified several circadian loci with evidence of adaptive introgression or latitudinal clines in allele frequency.

**Conclusions:** These findings identify differences in circadian gene regulation between modern humans and archaic hominins and support the contribution of introgression via coordinated effects on variation in human chronotype.

**SIGNIFICANCE STATEMENT:** Interbreeding between humans and Neanderthals created the potential for adaptive introgression as humans moved into environments that had been populated by Neanderthals for hundreds of thousands of years. Here we discover lineage-specific genetic differences in circadian genes and their regulatory elements between humans and Neanderthals. We show that introgressed alleles are enriched for effects on circadian gene regulation, consistently increase propensity for morningness in Europeans, and show evidence of adaptive introgression or associations between latitude and frequency. These results expand our understanding of how the genomes of humans and our closest relatives responded to environments with different light/dark cycles, and demonstrate a coordinated contribution of admixture to human chronotype in a direction that is consistent with adaptation to higher latitudes.

## INTRODUCTION

All anatomically modern humans (AMH) trace their origin to the African continent around 300 thousand years ago (ka) (Stringer, 2016; Hublin *et al*., 2017), where environmental factors shaped many of their biological features. Approximately seventy-thousand years ago (Bae, Douka, and Petraglia 2017), the ancestors of modern Eurasian AMH began to migrate out of Africa, where they were exposed to diverse new environments. In Eurasia, the novel environmental factors included greater seasonal variation in temperature and photoperiod.

Changes in the pattern and level of light exposure have biological and behavioral consequences in organisms. For example, *D. melanogaster* that are native to Europe harbor a polymorphism in *timeless*, a key gene in the light response of the circadian system, that follows a latitudinal cline in allele frequency (Sandrelli et al. 2007; Tauber et al. 2007). The ancestral haplotype produces a short TIM (S-TIM) protein that is sensitive to degradation by light because of its strong affinity to cryptochromes (CRY), photoreceptor proteins involved in the entrainment of the circadian clock. An insertion of a G nucleotide in the 5’ coding region of the gene originated approximately 10 kya in Europe and created a start codon that produces a new long TIM isoform (L-TIM). The L-TIM variant has a lower affinity to CRY, creating a change in photosensitivity and altering the length of the period. L-TIM flies are at a higher frequency in southern Europe, while S-TIM flies are more prevalent in northern Europe. Another example is found in pacific salmon. Chinook salmon (*Oncorhynchus tshawytscha*) populations show a latitudinal cline in the frequency and length of repeat motifs in the gene *OtsClock1b*, strongly suggesting that this locus is under selection associated with latitude and photoperiod (O’Malley, Ford, and Hard 2010; O’Malley and Banks 2008). The evolution of circadian adaptation to diverse environments has also been widely studied in insects, plants (Michael *et al*., 2003; Zhang *et al*., 2008), and fishes, but it is understudied in humans. Adaptive processes could have helped to align human biology and chronotype to new natural conditions.

Previous studies in humans found a correlation between latitude and chronotype (morningness vs. eveningness) variation (Leocadio-Miguel et al. 2017; Lowden et al. 2018; Randler and Rahafar 2017) and a latitudinal cline in some circadian allele frequencies (Dorokhov *et al*., 2018; Putilov, Dorokhov and Poluektov, 2018; Putilov *et al*., 2019), highlighting the contribution of the environment to behavior and circadian biology. Many human health effects are linked to the misalignment of chronotype (Knutson and von Schantz 2018), including cancer, obesity (Gyarmati *et al*., 2016; Papantoniou *et al*., 2016, 2017; Gan *et al*., 2018; Shi *et al*., 2020; Yousef *et al*., 2020), and diabetes (Gan *et al*., 2015; Larcher *et al*., 2015, 2016). There is also evidence of a correlation between evening chronotype and mood disorders, most notably seasonal affective disorder (SAD), depression, and worsening of bipolar disorder episodes (Srinivasan *et al*., 2006; Kivelä, Papadopoulos and Antypa, 2018; Taylor and Hasler, 2018).

Thus, we hypothesize that the differences in geography and environment encountered by early AMH populations moving into higher latitudes created potential for circadian misalignment and health risk.

Although AMHs arrived in Eurasia ∼70 ka, other hominins (e.g., Neanderthals and Denisovans) lived there for more than 400 ka (Arnold *et al*., 2014; Meyer *et al*., 2014, 2016). These archaic hominins diverged from AMHs around 700 ka (Meyer *et al*., 2012; Prüfer *et al*., 2014, 2017; Nielsen *et al*., 2017; Gómez-Robles, 2019; Mafessoni *et al*., 2020), and as a result, the ancestors of AMHs and archaic hominins evolved under different environmental conditions. While there was substantial variation in the latitudinal ranges of each group, the Eurasian hominins largely lived at consistently higher latitudes and, thus, were exposed to higher amplitude seasonal variation in photoperiods. Given the influence of environmental cues on circadian biology, we hypothesized that these separate evolutionary histories produced differences in circadian traits adapted to the distinct environments.

When AMH migrated into Eurasia, they interbred with the archaic hominins that were native to the continent, initially with Neanderthals (Green et al. 2010; Villanea and Schraiber 2019) around 60 ka (Sankararaman et al. 2012; Skoglund and Mathieson 2018) and later with Denisovans (Jacobs et al. 2019). Due to this, a substantial fraction (>40%) of the archaic variation remains in present-day Eurasians (Skov et al. 2020; Vernot and Akey 2014), although each human individual carries only ∼2% DNA of archaic ancestry (Vernot *et al*., 2016; Prüfer *et al*., 2017). Most of the archaic ancestry in AMH was subject to strong negative selection, but some of these introgressed alleles remaining in AMH populations show evidence of adaptation (Racimo *et al*., 2015; Gower *et al*., 2021). For example, archaic alleles have been associated with differences in hemoglobin levels at higher altitude in Tibetans, immune resistance to new pathogens, levels of skin pigmentation, and fat composition (Huerta-Sánchez *et al*., 2014; Racimo *et al*., 2015, 2017; Dannemann and Kelso, 2017; Racimo, Marnetto and Huerta-Sánchez, 2017; McArthur, Rinker and Capra, 2021). Previous work also suggests that introgressed alleles could have adaptively influenced human chronotype. First, a phenome-wide association study (PheWAS) in the UK Biobank found loci near *ASB1* and *EXOC6* with introgressed variants that significantly associated with self-reported sleeping patterns (Dannemann and Kelso, 2017). One of these alleles showed a significant association between frequency and latitude. Second, summarizing effects genome-wide, introgressed alleles are also moderately enriched for heritability of chronotype compared to non-introgressed alleles (McArthur, Rinker and Capra, 2021). These results suggest a potential role for introgressed alleles in adaptation to pressures stemming from migration to higher latitudes.

Motivated by the potential for a role of archaic introgression in AMH circadian variation, we explore two related questions: 1) Can comparative genomic analysis identify differences in AMH and archaic hominin circadian biology?, and 2) Do introgressed archaic alleles influence human circadian biology? Understanding the ancient history and evolution of chronotypes in humans will shed light on human adaptation to high latitudes and provide context for the genetic basis for the modern misalignment caused by the development of technology and night shiftwork.

## RESULTS

### Did archaic hominins and modern humans diverge in circadian biology?

Following divergence ∼700,000 years ago (ka) (Nielsen *et al*., 2017; Gómez-Robles, 2019), archaic hominins and AMH were geographically isolated, resulting in the accumulation of lineage-specific genetic variation and phenotypes (Figure 1). In the next several sections, we evaluate the genomic evidence for divergence in circadian biology between archaic hominin and modern human genomes.

**Figure 1.**
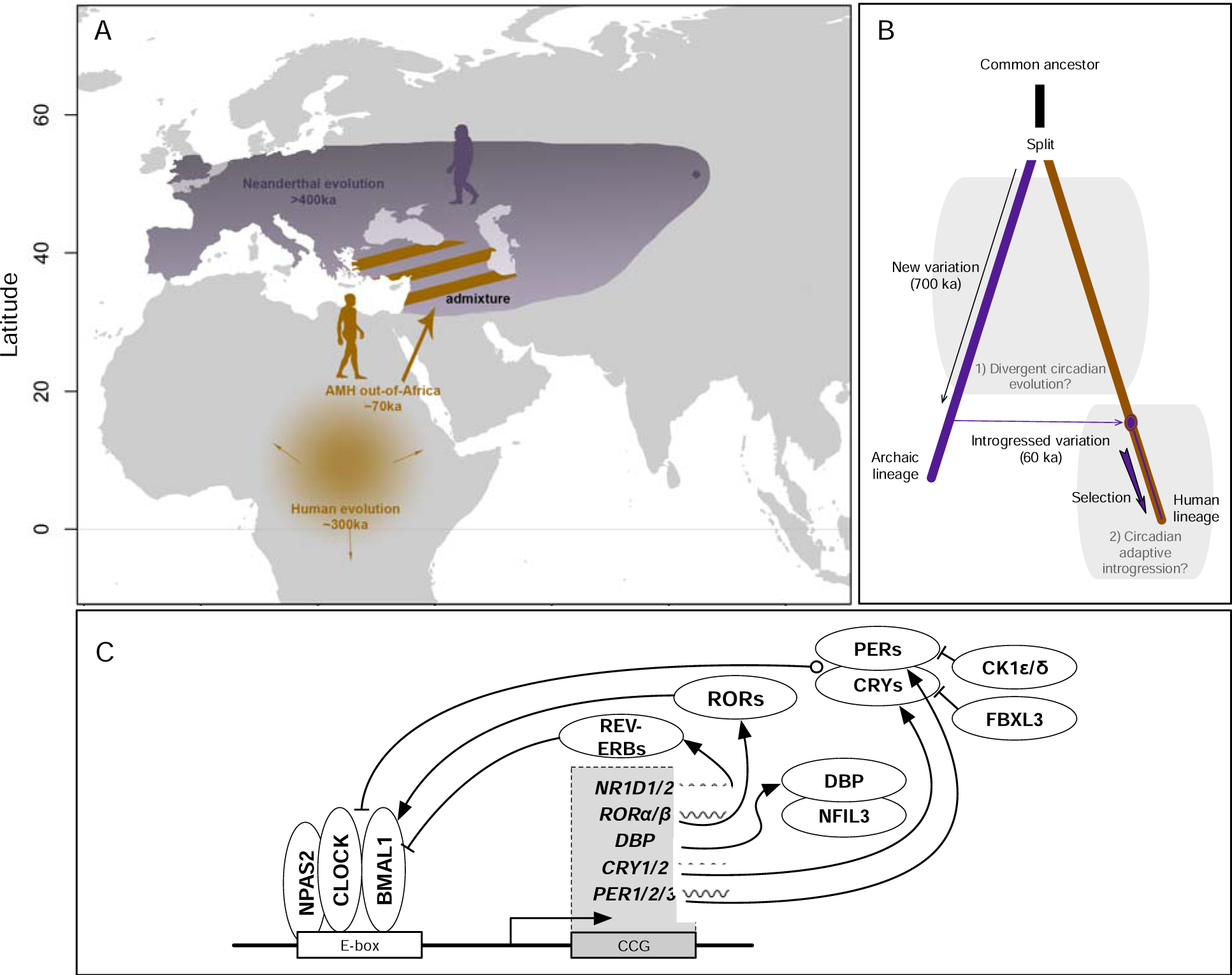
Did the sharing of functionally diverged alleles from archaic hominins influence human circadian biology? A) Anatomically modern humans and archaic hominins evolved separately at different latitudes for hundreds of thousands of years. The ancestors of modern Eurasian humans left Africa approximately 70 thousand years ago (ka) and admixed with archaics, likely in southwestern Asia. The shaded purple range represents the approximate Neanderthal range. The purple dot represents the location of the sequenced Denisovan individual in the Altai Mountains; the full range of Denisovans is currently unknown. Silhouettes from phylopic.org. **B)** After the split between the human and archaic lineages, each group accumulated variation and evolved in their respective environments for approximately 700 ka. We first test for evidence for divergent circadian evolution during this time. Humans acquired introgressed alleles from Neanderthals and from Denisovans around 60 and 45 ka, respectively. These alleles experienced strong selective pressures; however, ∼40% of the genome retains archaic ancestry in some modern populations. The second question we explore is whether introgression made contributions to human circadian biology. **C)** The core circadian clock machinery is composed of several transcription factors (ovals) that dimerize and interact with E-box enhancer elements and each other to create a negative feedback loop. We defined a set of 246 circadian genes through a combination of literature search, expert knowledge, and existing annotations (Table S1; Figure S1; Methods). Lines with arrows represent activation, and lines with bars represent suppression.

### Identifying archaic-hominin-specific circadian gene variation

With the sequencing of several genomes of archaic hominins, we now have a growing, but incomplete, catalog of genetic differences specific to modern and archaic lineages. Following recent work (Kuhlwilm and Boeckx, 2019), we defined archaic-specific variants as genomic positions where archaic hominins (Altai Neanderthal, Vindija Neanderthal, and Denisovan) all have the derived allele while in humans the derived allele is absent or present at such an extremely low frequency in the 1000 Genome Project (<0.00001) that it is likely an independent occurrence. We defined human-specific variants as positions where all individuals in the 1000 Genomes Project carry the derived allele and all the archaics carry the ancestral allele.

We evaluated archaic-specific variants for their ability to influence proteins, splicing, and regulation of 246 circadian genes (Methods). The circadian genes were identified by a combination of literature search, expert knowledge, and existing annotations (Table S1; Figure S1; Methods). The core circadian clock machinery is composed of a dimer between the CLOCK and ARNTL (BMAL1) transcription factors, which binds to E-box enhancer elements and activates the expression of the Period (*PER1/2/3*) and Cryptochrome (*CRY1/2*) genes (Figure 1C). PERs and CRYs form heterodimers that inhibit the positive drive of the CLOCK-BMAL1 dimer on E-boxes, inhibiting their own transcription in a negative feedback loop. CLOCK- BMAL1 also drives the expression of many other clock-controlled genes (CCG), including *NR1D1/2* (Nuclear Receptor Subfamily 1 Group D Member 1 and 2), *RORA/B* (RAR Related Orphan Receptor A and B), and *DBP* (D-Box Binding PAR BZIP Transcription Factor). ROR and REV-ERB are transcriptional regulators of BMAL1. CK1 binds to the PER/CRY heterodimer, phosphorylating PER and regulating its degradation. Similarly, FBXL3 marks CRY for degradation. Beyond the core clock genes, we included other upstream and downstream genes that are involved in maintenance and response of the clock (Table S1; Figure S1).

We identified 1,136 archaic-specific variants in circadian genes, promoters, and candidate distal cis-regulatory elements (cCREs). The circadian genes with the most archaic- specific variants are *CLDN4*, *NAMPT*, *LRPPRC*, *ATF4*, and *AHCY* (125, 112, 110, 104, 102 respectively) (Table S2).

### Fixed human- and archaic-specific variants are enriched in circadian genes and associated regulatory elements

After the archaic and AMH lineages diverged, each group accumulated genetic variation specific to each group. Variants fixed in each lineage are likely to be enriched in genomic regions that influence traits that experienced positive selection. We tested whether human- and archaic-specific fixed variants are enriched compared to other variants in circadian genes, their promoters, and in annotated candidate cis-regulatory elements within 1 megabase (Mb) (Figure 2). We found that human- and archaic-specific fixed variants are enriched in circadian genes (Fisher’s exact test; human: OR=1.84, P=7.06e-12; archaic: OR=1.13, P=0.023) and distal regulatory elements (Fisher’s exact test; human: OR=1.25, P=8.39e-4; archaic: OR=1.16, P=6.15e-5) compared to variants derived on each lineage, but not fixed. Promoter regions have a similar enrichment pattern as that in gene and regulatory regions, but the p-values are high (Fisher’s exact test; human: OR=1.21, P=0.65; archaic: OR=1.09, P=0.63). This is likely due to the small number of such variants in promoters. These results suggest that both groups had a greater divergence in genomic regions related to circadian biology than expected.

**Figure 2.**
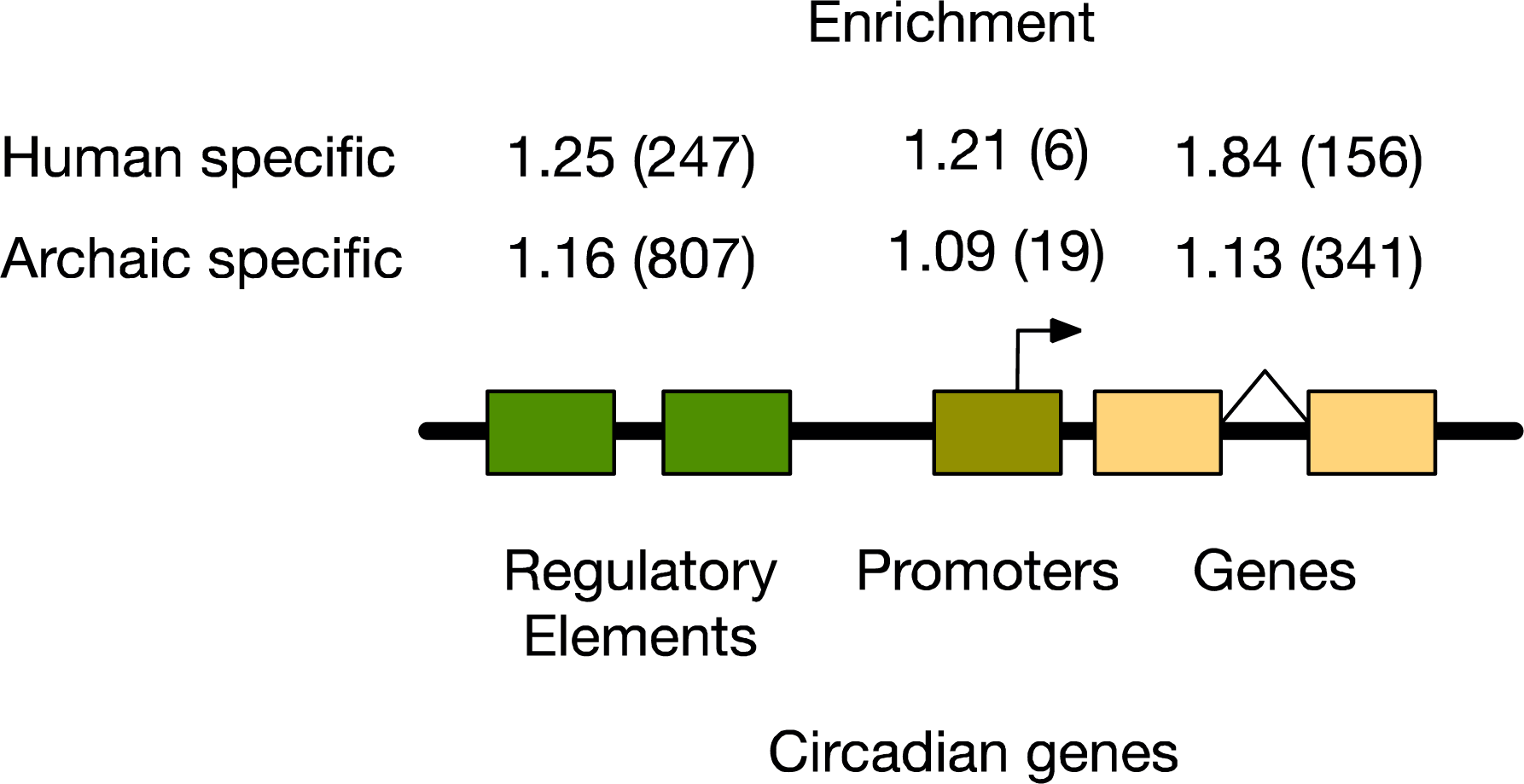
Human- and archaic-specific fixed variants are enriched in circadian regulatory, promoter, and gene regions. Human-specific fixed variants are significantly enriched compared to variants that are not fixed in circadian regulatory elements (Fisher’s exact: OR=1.25, P=8.39e- 4) and gene regions (Fisher’s exact: OR=1.84, P=7.06e-12). Promoters show a similar enrichment, but the higher p-value is the result of the small number of variants (Fisher’s exact test: OR=1.21, P=0.65). Likewise, archaic-specific variants are enriched in circadian regulatory regions (Fisher’s exact: OR=1.16, P=6.15e-5) and gene regions (Fisher’s exact: OR=1.13, P=0.023), with the promoters showing a similar trend (Fisher’s exact test: OR=1.09, P=0.63).

The numbers in parentheses give the counts of fixed variants observed in each type of element. Regulatory elements were defined based on the ENCODE candidate cis-regulatory elements.

### Several core circadian genes have evidence of alternative splicing between humans and archaic hominins

We find only two archaic-specific coding variants in circadian genes: one missense and one synonymous. The missense variant (hg19: chr17_46923411_A_G) is in the gene *CALCOCO2*, calcium-binding and coiled-coil domain-containing protein 2. SIFT, PolyPhen, and CADD all predict that the variant does not have damaging effects. The second variant (hg19: chr7_119914770_G_T) is in the gene *KCND2*, which encodes a component of a voltage-gated potassium channel that contributes to the regulation of the circadian rhythm of action potential firing, but it is synonymous and the variant effect predictors suggest it is tolerated.

To explore potential splicing differences in circadian genes between humans and archaics, we applied SpliceAI to predict whether any sequence differences between modern humans and archaics are likely to modify splicing patterns. Four archaic individuals were included in this analysis (the Altai, the Vindija, the Chagyrskaya Neanderthals, and the Altai Denisovan) (Meyer *et al*., 2012; Prüfer *et al*., 2014, 2017; Mafessoni *et al*., 2020). We found that 28 genes contained at least one archaic-specific variant predicted to result in alternative splicing in archaics. These included several of the core clock genes *CLOCK*, *PER2*, *RORB*, *RORC*, and *FBXL13* (Figure 3A,C; Table S3). For example, the variant chr2:239187088-239187089 in the 1st intron of *PER2* is predicted to result in a longer 5’ UTR. The splice-altering variants were largely specific to the two different archaic linages (Figure 3A), with 13 specific to the Denisovan, 8 shared among the three Neanderthals, and only one shared among all four archaic individuals.

**Figure 3.**
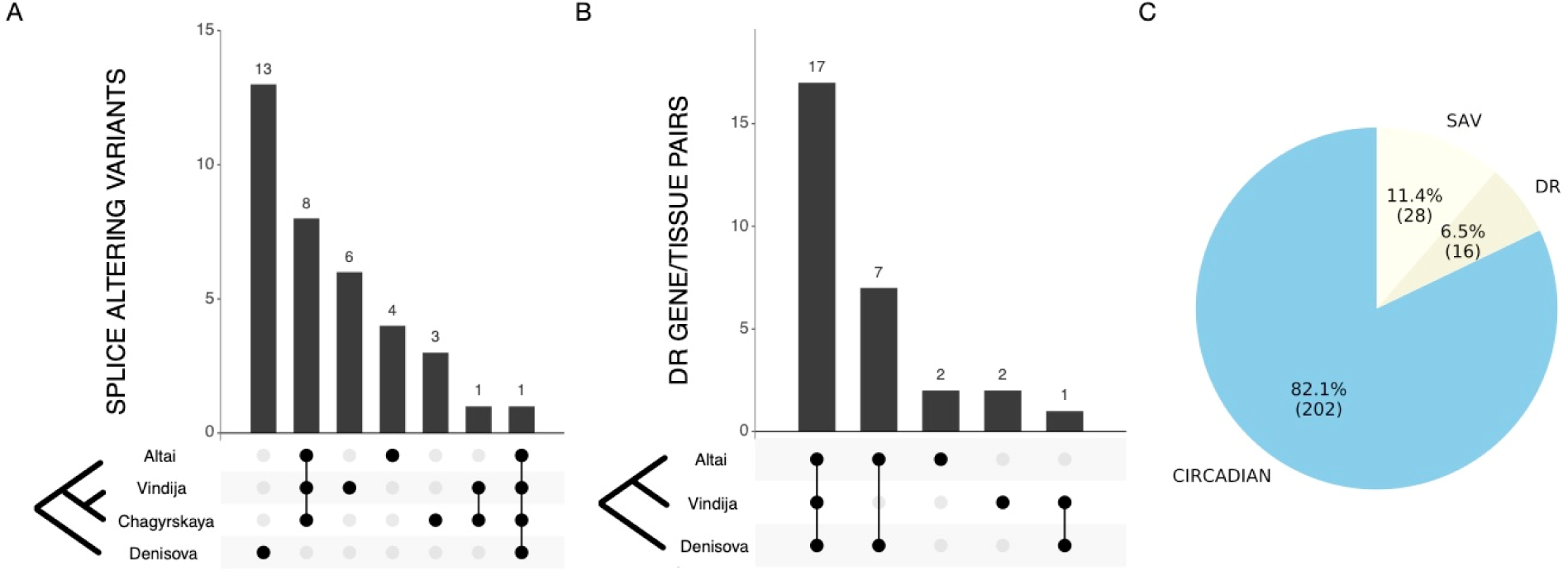
Many circadian genes have evidence of alternative splicing and divergent regulation between modern and archaic hominins. A) The distribution of the 28 predicted archaic-specific splice-altering variants (SAV) in circadian genes across archaic individuals. Most are specific to either the Denisovan or Neanderthal lineage (Table S3). **B)** The sharing of predicted divergently regulated (DR) gene/tissue pairs across three archaic individuals. (Predictions were not available for the Chagyrskaya Neanderthal.) Seventeen divergently regulated gene/tissue pairs were present in all three archaics (representing 16 unique genes).

### Circadian gene regulatory divergence between humans and archaic hominins

Given the enrichment of variants in regulatory regions of circadian genes, we sought to explore the potential for differences in circadian gene regulation between humans and archaics with causes beyond single lineage-specific variants. We leveraged an approach we recently developed for predicting gene regulatory differences between modern and archaic individuals from combinations of genetic variants (Colbran *et al*., 2019). The approach uses PrediXcan, an elastic net regression method, to impute gene transcript levels in specific tissues from genetic variation. Previous work demonstrated that this approach has a modest decrease in performance when applied to Neanderthals, but that it can accurately applied between humans and Neanderthals for thousands of genes. Here, we quantify differences in predicted regulation of the 246 circadian genes between 2,504 humans in the 1000 Genomes Project (1000 Genomes Project Consortium, 2010) and the archaic hominins. The predicted regulation values are normalized to the distribution in the training set from the Genotype Tissue Expression Atlas (GTEx).

Additionally, 7 gene/tissue DR pairs are shared between the Altai Neanderthal and the Denisovan individual. One pair is shared between the Vindija Neanderthal and the Denisovan (Table S4). **C)** The proportion of circadian genes containing archaic splice-altering variants predicted by SpliceAI (SAV; 11.4%) or divergently regulated circadian genes predicted by PrediXcan (DR; 6.5%). Thus, 17.9% of the circadian genes are predicted to contain differences to AMH via these mechanisms.

We first analyzed gene regulation predictions in the core circadian clock genes. Archaic gene regulation was at the extremes of the human distribution for many core clock genes including *PER2*, *CRY1*, *NPAS2*, *RORA, NR1D1* (Figure 4; Figure S2). For example, the regulation of *PER2* in the Altai and Vindija Neanderthals is lower than 2,491 of the 2,504 (99.48%) modern humans considered. The Denisovan has a predicted *PER2* regulation that is lower than 2,410 (96.25%). Expanding to all circadian genes and requiring archaic regulation to be more extreme than all humans (Methods), we identified 24 circadian genes across 23 tissues with strong divergent regulation between humans and at least one archaic hominin (Figure 3B; Table S4). For example, all archaic regulation values for *RORA*, a core clock gene, are lower than for any of the 2,504 modern humans. We found that 16 of these genes (Figure S3; Table S4), including *RORA*, *MYBBP1A*, and *TIMELESS*, were divergently regulated in all archaic individuals. This represents 6.5% of all the circadian genes (Figure 3C). Surprisingly, the two Neanderthals only shared one DR gene not found in the Denisovan, while the Altai Neanderthal and Denisovan shared seven not found in Vindija (Figure 3B). The Altai and Vindija Neanderthals represent deeply diverging lineages, and this result suggests that they may have experienced different patterns of divergence in the regulation of their circadian genes.

**Figure 4.**
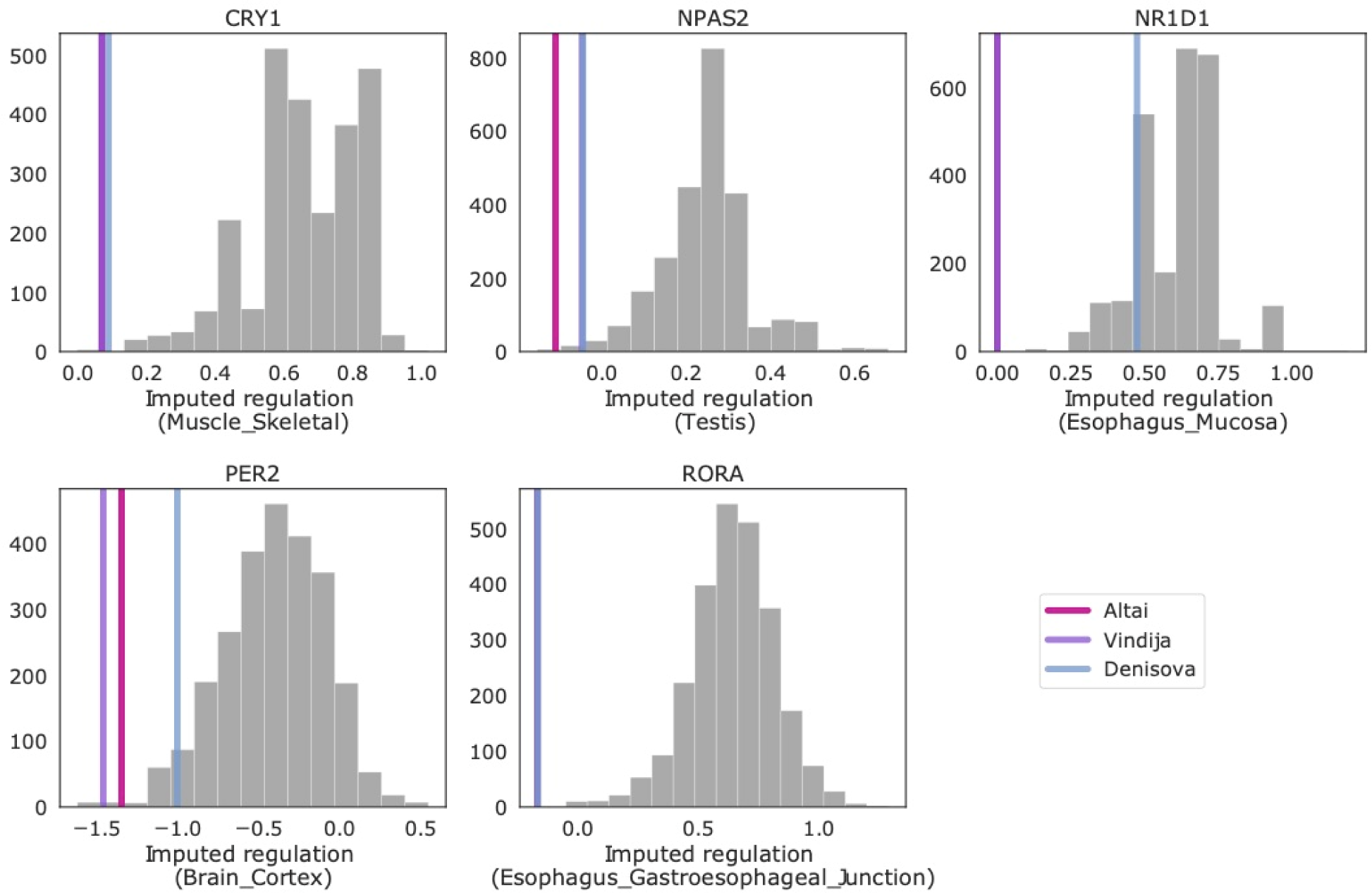
Many circadian genes are divergently regulated between modern humans and archaic hominins. Comparison of the imputed regulation of core circadian genes between 2504 humans in 1000 Genomes Phase 3 (gray bars) and three archaic individuals (vertical lines). For each core circadian gene, the tissue with the lowest average P-value for archaic difference from humans is plotted. Archaic gene regulation is at the extremes of the human distribution for several core genes: *CRY1*, *PER2*, *NPAS2, NR1D1 RORA*. See Figure S2 for all core clock genes and Figure S3 for all divergently regulated circadian genes.

Given these differences in circadian gene regulation between humans and archaics, we tested whether circadian genes are more likely to be divergently regulated than other gene sets. Each archaic individual shows nominal enrichment for divergent regulation of circadian genes, and the enrichment was stronger (∼1.2x) in the Altai Neanderthal and Denisovan individual. However, given the small sample size, the P-values are moderate (Permutation test; Altai: OR=1.21, P=0.19, Vindija: OR=1.05, P=0.43, Denisovan: OR=1.20, P=0.24).

### Did introgressed archaic variants influence modern human circadian biology?

The previous sections demonstrate lineage-specific genetic variation in many genes and regulatory elements essential to the function of the core circadian clock and related pathways. Given this evidence of functional differences between archaic hominins and AMH in these systems, we next evaluated the influence of archaic introgression on AMH circadian biology.

### Introgressed variants are enriched in circadian gene eQTL

Given the differences between archaic and modern sequences of circadian genes and their regulatory elements, we investigated whether Neanderthal introgression contributed functional circadian variants to modern Eurasian populations. We considered a set of 863,539 variants with evidence of being introgressed from archaic hominins to AMH (Browning *et al*., 2018). These variants were identified using the Sprime algorithm, which searches for regions containing a high density of alleles in common with Neanderthals and not present or at very low frequency in Africans. Since many approaches have been developed to identify introgressed variants, we also considered two stricter sets: 47,055 variants that were supported by all of six different introgression maps (Sankararaman *et al*., 2014; Vernot *et al*., 2016; Browning *et al*., 2018; Steinrücken *et al*., 2018; Skov *et al*., 2020; Schaefer, Shapiro and Green, 2021) and 755,653 variants that were supported by Sprime and at least one other introgression map. As described below, our main results replicated on both of these stricter sets.

We first tested whether the presence of introgressed variants across modern individuals associated with the expression levels of any circadian genes, i.e., whether the introgressed variants are expression quantitative trait loci (eQTL). We identified 3,857 introgressed variants associated with the regulation of circadian genes in modern non-Africans (Table S5). The genes *PTPRJ*, *HTR1B*, *NR1D2*, *CLOCK*, and *ATOH7* had the most eQTL (304, 273, 262, 256, and 252respectively). We found introgressed circadian eQTL for genes expressed in all tissues in GTEx, except kidney cortex. Notably, several of these circadian genes (e.g., *NR1D2* and *CLOCK*) with introgressed eQTL were also found to be divergently regulated in our comparison of modern and archaic gene regulation. This indicates that some of the archaic-derived variants that contributed to divergent regulation were retained after introgression and continue to influence circadian regulation in modern humans.

Introgressed variants are significantly more likely to be eQTL for circadian genes than expected by chance from comparison to all eQTL (Figure 5A; Fisher’s exact test: OR=1.45, P=9.71e-101). The stricter set of introgressed variants identified by Browning et al. plus at least one other introgression map had similar levels of eQTL enrichment for circadian genes (OR=1.47; P=2.4e-103). The highest confidence set of introgressed variants that were identified by all six maps considered had even stronger enrichment (OR=1.68; P=6.5e-23).

**Figure 5.**
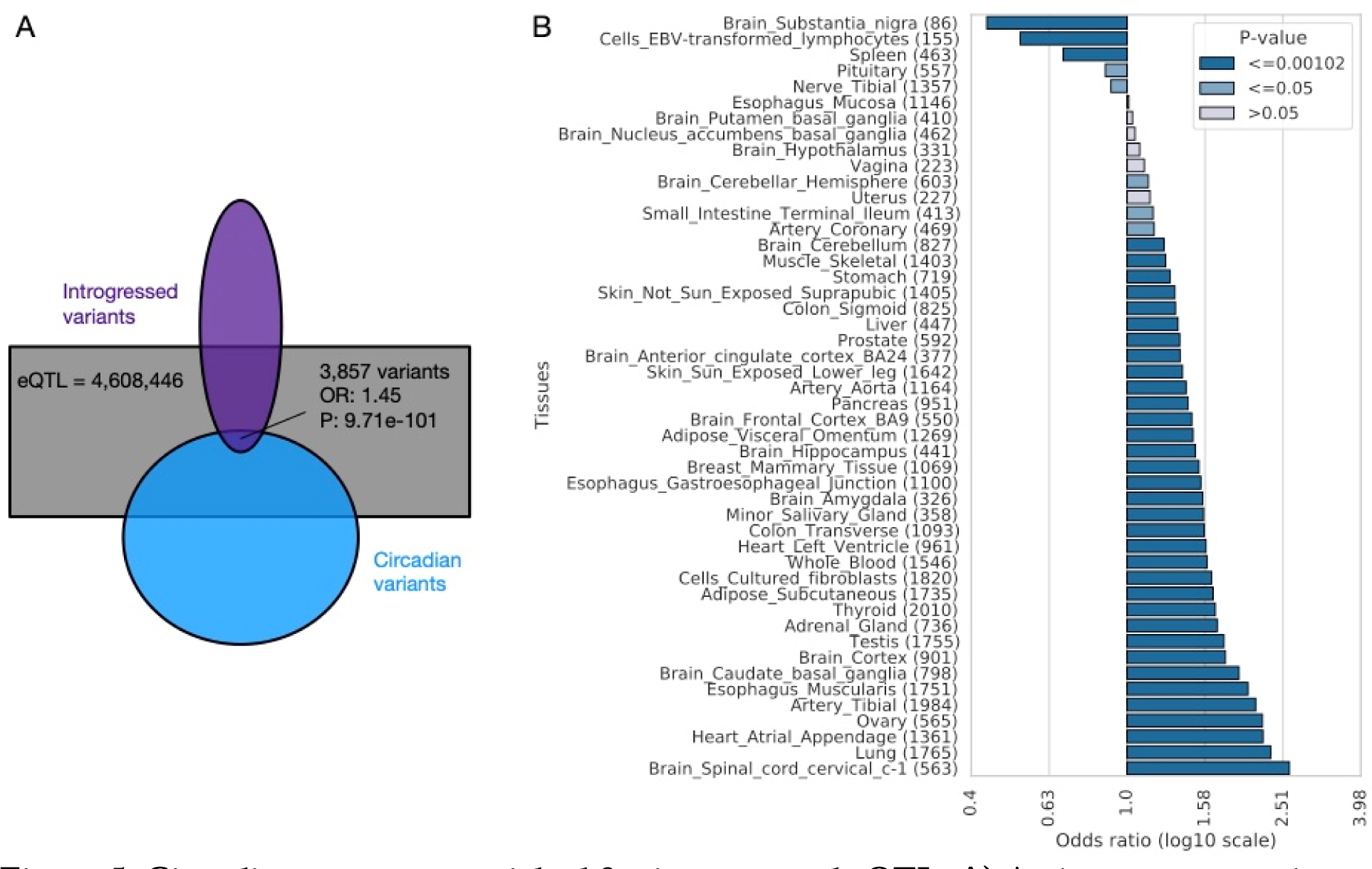
Circadian genes are enriched for introgressed eQTL. A) Archaic introgressed variants are more likely to be eQTL for circadian genes in GTEx than for non-circadian genes (Fisher’s exact test: OR=1.45, P=9.71e-101). Purple represents the set of introgressed variants, and blue represents the set of circadian variants. 3,857 are introgressed eQTL in circadian genes. Gray represents the universe of all GTEx eQTLs lifted over to hg19. The overlaps are not to scale. **B)** The enrichment for circadian genes among the targets of introgressed eQTLs in each GTEx tissue. Introgressed eQTL in most tissues show significant enrichment for circadian genes (Fisher’s exact test; Table S7). Kidney cortex did not have any circadian introgressed eQTLs and thus is not shown. Numbers inside the parenthesis indicate the count of variants in each tissue.

Most core circadian genes are expressed broadly across tissues; the fraction expressed in each GTEx tissue ranges from 57% (whole blood) to 83% in testis, and an average of 72% (Table S6). As a result, we anticipated that the enrichment of introgressed variants among eQTL for circadian genes would hold across tissues. Examining the associations in each tissue, we found that introgressed eQTL showed significant enrichment for circadian genes in most tissues (34 of 49; Figure 5B; Table S7) and trended this way in all but five. Given that tissues in GTEx have substantial differences in sample size and cellular heterogeneity, statistical power to detect enrichment differs. We anticipate that this is the main driver of differences in enrichment across tissues.

These results suggest that circadian pressures were widespread across tissues. Given the previously observed depletion for introgressed variants in regulatory elements and eQTL (Petr *et al*., 2019; Rinker *et al*., 2020; Telis, Aguilar and Harris, 2020), this enrichment for circadian genes among introgressed eQTL is surprising and suggests that the archaic circadian alleles could have been beneficial after introgression.

Gray bars indicate lack of statistical significance; light blue bars indicate nominal significance (p<= 0.05); and dark blue bars indicate significance at the 0.05 level after Bonferroni multiple testing correction (p <= 0.00102).

### Introgressed variants predominantly increase propensity for morningness

After observing that circadian gene expression is influenced by archaic variants, we evaluated whether these effects are likely to result in a change in organism-level phenotype. To do this, we evaluated evidence that introgressed variants influence chronotype. The heritability of chronotype has been estimated in a range from 12 to 38% (Jones *et al*., 2016, 2019; Lane *et al*., 2016), and previous studies have identified two introgressed loci associated with sleep patterns (Dannemann and Kelso, 2017; Putilov *et al*., 2019). We recently found modest enrichment for heritability of chronotype (morning/evening person phenotype in a GWAS of the UK Biobank) among introgressed variants genome-wide using stratified LD score regression (heritability enrichment: 1.58, P=0.25) (McArthur, Rinker and Capra, 2021). This analysis also suggested that introgressed variants were more likely to increase morningness.

To test for this proposed directional effect, we calculated the cumulative fraction of introgressed loci associated with chronotype in the UK Biobank that increase morningness (after collapsing based on LD at R^2^>0.5 in EUR). The introgressed loci most strongly associated with chronotype increase propensity for morningness (Figure 6; Table S8; Table S9). As the strength of the association with morningness decreases, the bias begins to decrease, but the effect is maintained well past the genome-wide significance threshold (P<5e-8). When focusing the analysis on introgressed variants in proximity (<1 Mb) to circadian genes, the pattern becomes even stronger. The bias toward morningness remains above 80% at the genome-wide significance threshold. This result also held when limiting to introgressed variants found in Browning plus one or all other introgression maps considered (Figure S4). This suggests that introgressed variants act in a consistent direction on chronotype, especially when they influence circadian genes.

**Figure 6.**
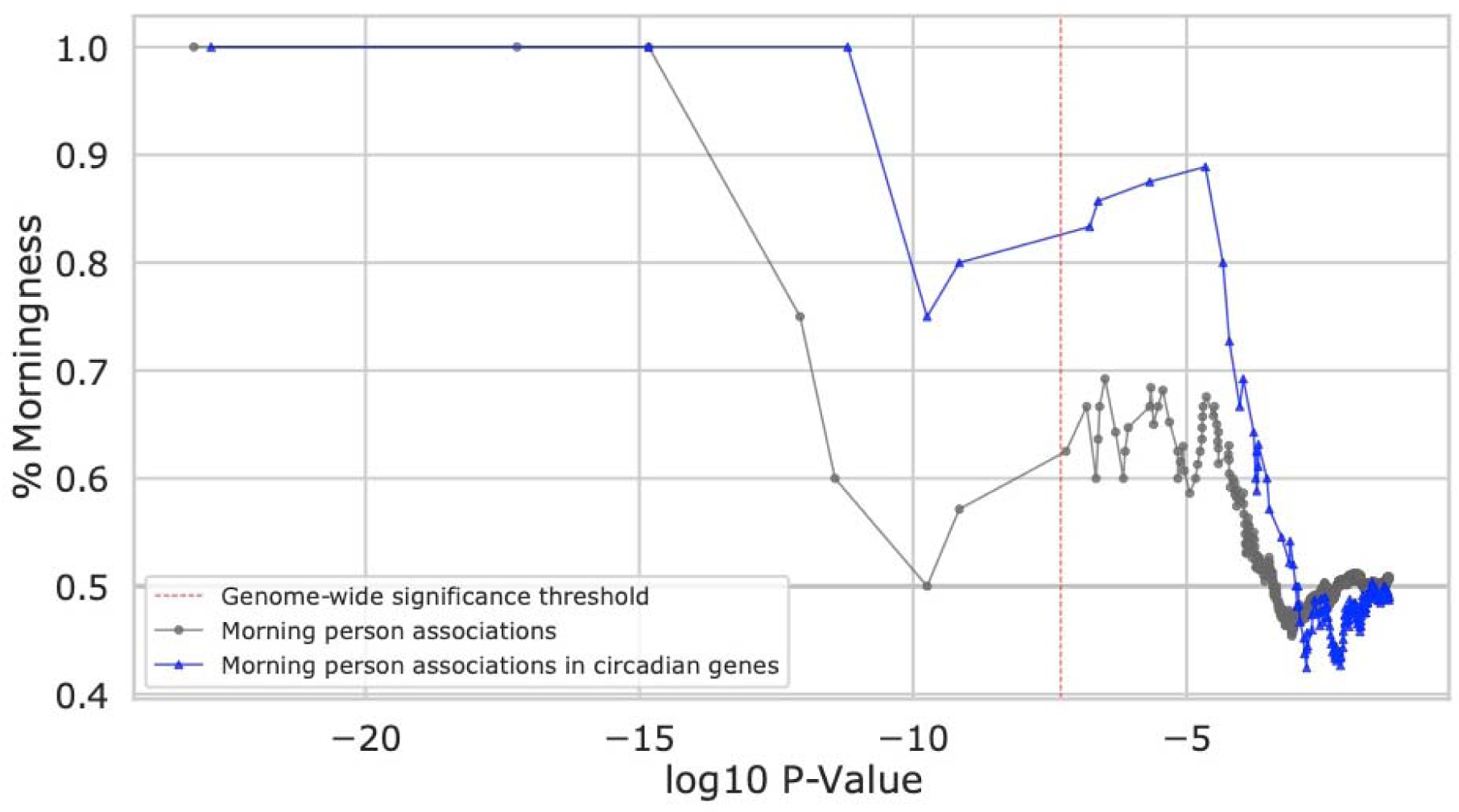
Introgressed variants associate with increased morningness. The cumulative fraction of introgressed loci significantly associated with the morning vs. evening person trait in the UK Biobank that increase morningness (y-axis) at a given p-value threshold (x-axis).

Introgressed loci associated with chronotype are biased towards increasing morningness, and this effect is greatest at the most strongly associated loci. Introgressed variants nearby (<1 Mb) circadian genes (blue) are even more strongly biased towards increasing morningness than introgressed variants overall (gray). Each dot (triangle) represents an associated locus; variants were clumped by LD for each set (R^2^>0.5 in EUR).

Circadian rhythms are involved in a wide variety of biological systems. To explore other phenotypes potentially influenced by the introgressed circadian variants, we evaluated evidence for pleiotropic associations. First, we retrieved all the genome-wide associations reported for introgressed variants in the Open Targets Genetics (https://genetics.opentargets.org) database, which combines GWAS data from the GWAS Catalog, UK Biobank, and several other sources.

Introgressed circadian variants are associated with traits from a diverse range of categories (Table S10). Associations with blood related traits are by far the most common; however, this is likely because they have more power in the UK Biobank. Overall, circadian introgressed variants are significantly more likely to have at least one trait association than introgressed variants not in the circadian set (Fisher’s exact test: OR=1.25, P=7.03e-25) (Figure S5A). The circadian variants also associate with significantly more traits per variant than the non-circadian set (Mann-Whitney U: P=9.93e-14) (Figure S5B; Table S11). These results suggest effects for introgressed circadian variants beyond chronotype.

### Evidence for adaptive introgression at circadian loci

The gene flow from Eurasian archaic hominins into AMH contributed to adaptations to some of the new environmental conditions encountered outside of Africa (Racimo *et al*., 2015). The above analyses demonstrate the effects of introgressed variants on circadian gene regulation and chronotype. To explore whether these circadian regions show evidence of adaptive introgression, we considered three sets of introgressed regions predicted to have contributed to AMH adaptation: one from an outlier approach based on allele frequency statistics (Racimo, Marnetto and Huerta-Sánchez, 2017) and two from recent machine learning algorithms: *genomatnn* (Gower *et al*., 2021) and *MaLAdapt* (Zhang *et al*., 2023). We intersected the circadian introgressed variants with the adaptive introgression regions from each method.

We identified 47 circadian genes with evidence of adaptive introgression at a nearby variant from at least one of the methods (Table S12). No region was supported by all three methods; however, six were shared between Racimo and *MaLAdapt* and three were shared by Racimo and *genomatnn*. The relatively small overlap between these sets underscores the challenges of identifying adaptive introgression. Nonetheless, these represent promising candidate regions for further exploration of the effects of introgressed variants on specific aspects of circadian biology. For example, an introgressed haplotype on chr10 tagged by rs76647913 was identified by both *MaLAdapt* and Racimo. This introgressed haploype is an eQTL for the nearby *ATOH7* gene in many GTEx tissues. *ATOH7* is a circadian gene that is involved in retinal ganglion cell development, and mice with this gene knocked out are unable to entrain their circadian clock based on light stimuli (Brzezinski *et al*., 2005).

### Latitudinal clines for introgressed circadian loci

Motivated by the previous discovery of an introgressed haplotype on chr2 that is associated with chronotype and increases in frequency with latitude (Dannemann and Kelso, 2017; Putilov *et al*., 2019), we also tested each introgressed circadian variant for a correlation between allele frequency and latitude in modern non-African populations from the 1000 Genomes Project.

The strongest association between latitude and frequency was a large chromosome 2 haplotype that contains the previously discovered introgressed SNP (rs75804782, R=0.85) associated with chronotype. This haplotype is present in all non-African populations, and rs61332075 showed the strongest latitudinal cline (R=0.87). The second strongest consisted of a smaller haplotype of introgressed variants a few kb upstream of the previous haplotype (tagged by rs35333999 and rs960783) that overlaps the core circadian gene *PER2*. These variants have a correlation between latitude and frequency of ∼0.68 They are also in moderate LD (R^2^ of ∼0.35 in EUR) with an additional introgressed variant (rs62194932) that has a similar latitudinal cline of 0.70 (Figure S6; Table S13). These variants are each in very low LD with the previously discovered haplotype (R^2^ of ∼0.01) and are each supported by multiple introgression maps.

Moreover, these introgressed variants are absent in all EAS populations, absent or at very low frequency in SAS (<3%), and at higher frequency in EUR populations (∼13%).

The EUR-specific introgressed variant rs35333999 causes a missense change in the PER2 protein (V903I) that overlaps a predicted interaction interface with PPARG. PER2 controls lipid metabolism by directly repressing PPARG’s proadipogenic activity (Grimaldi *et al*., 2010). The rs62194932 variant is an eQTL of *HES6* in the blood in the eQTLGen cohort (Võsa *et al*., 2021). *HES6* encodes a protein that contributes to circadian regulation of LDLR and cholesterol homeostasis (Lee *et al*., 2012).

Thus, this genomic region, that includes circadian genes and introgressed variants associated with chronotype, has population-specific structure and at least two distinct sets of introgressed variants with latitudinal clines and functional links to lipid metabolism. *PER2* is also predicted to have lower gene regulation in archaic hominins than most humans (Figure 4). and the Vindija Neanderthal carries a lineage-specific variant in this gene that has splice-altering effects. These results together suggests that *PER2* may have experienced multiple functional changes in different modern and archaic lineages, with potential adaptive effects mediated by introgression.

We did not discover any other significant associations between latitude and frequency for other introgressed circadian loci. The rapid migration and geographic turnover of populations in recent human history is likely to obscure many latitude-dependent evolutionary signatures, so we did not anticipate many circadian loci would have a strong signal.

## DISCUSSION

The Eurasian environments where Neanderthals and Denisovans lived for several hundred thousand years are located at higher latitudes with more variable photoperiods than the landscape where AMH evolved before leaving Africa. Evaluating genetic variation that arose separately in each of the archaic and AMH lineages after their split ∼700 MYA, we identified lineage-specific genetic variation in circadian genes, their promoters, and flanking distal regulatory elements. We found that both archaic- and human-specific variants are observed more often than expected in each class of functional region. This result suggests that, while each group evolved separately during hundreds of thousands of years in divergent environments, both experienced pressure on circadian related variation. Leveraging sequence-based machine learning methods, we identified many archaic-specific variants likely to influence circadian gene splicing and regulation. For example, core clock genes (*CLOCK*, *PER2*, *RORB*, *RORC*, and *FBXL13*) have archaic variants predicted to cause alternative splicing compared to AMH. Several core genes were also predicted in archaics to be at the extremes of human gene regulation, including *PER2*, *CRY1*, *NPAS2*, *RORA, NR1D1*. Surprisingly, the Altai Neanderthal shared more divergent regulation in the circadian genes with the Denisovan individual than the Vindija Neanderthals. The two Neanderthals represent populations that were quite distantly diverged with substantially different histories and geographical ranges. The Denisovan and Altai Neanderthal also come from the same region in Siberia, while the Vindija Neanderthal came from a region in Croatia with slightly lower latitude.

Introgression introduced variation that first appeared in the archaic hominin lineage into Eurasian AMH. While most of this genetic variation experienced strong negative selection in AMH, a smaller portion is thought to have provided adaptive benefits in the new environments(Racimo *et al*., 2015). Given the divergence in many circadian genes’ regulation, we explored the landscape of introgression on circadian genes. We first looked at introgressed circadian variants that are likely to influence gene regulation in AMH. Variants in this set are observed more often than expected, suggesting the importance of maintaining circadian variation in the population. We also verified that these results held over variants identified by different methods for calling archaic introgression.

We then evaluated the association of these introgressed variants with variation in circadian phenotypes of Eurasians. We previously reported a modest enrichment among introgressed variants for heritability of the morning/evening person phenotype (McArthur, Rinker and Capra, 2021). Here, we further discovered a consistent directional effect of the introgressed circadian variants on chronotype. The strongest associated variants increase the probability of being a morning person in Eurasians.

While it is not immediately clear why increased morningness would be beneficial at higher latitudes, considering this directional effect in the context of clock gene regulation and the challenge of adaptation to higher latitudes suggests an answer. In present day humans, behavioral morningness is correlated with shortened period of the circadian molecular clockworks in individuals. This earlier alignment of sleep/wake with external timing cues is a consequence of a quickened pace of the circadian gene network (Brown *et al*., 2008). Therefore, the morningness directionality of introgressed circadian variants may indicate selection toward shortened circadian period in the archaic populations living at high latitudes. Supporting this interpretation, shortened circadian periods are required for synchronization to the extended summer photoperiods of high latitudes in *Drosophila*, and selection for shorter periods has resulted in latitudinal clines of decreasing period with increasing latitude, as well as earlier alignment of behavioral rhythms (Hut *et al*., 2013). In addition, *Drosophila* populations exhibit decreased amplitude of behavioral rhythms at higher latitudes which is also thought to aid in synchronization to long photoperiods (Hut *et al*., 2013).

Our finding that introgressed circadian variants generally decrease gene regulation of circadian genes suggests that they could lead to lower amplitude clock gene oscillations.

However, when assayed in present day humans there is not a strong correlation between the overall expression level of *NR1D1* and the transcriptional amplitudes of other clock genes within individuals (Brown *et al*., 2008), and quantitative modeling of the mammalian circadian clockworks suggests that stable clock gene rhythms can result across a wide range of absolute levels of gene expression as long as the stoichiometric ratios of key positive and negative clock genes are reasonably conserved (Kim and Forger, 2012). Interestingly, lower transcriptional amplitude of *NR1D1* does confer greater sensitivity of the present-day human clockworks to resetting stimuli, a potentially adaptive characteristic for high latitudes (Brown *et al*., 2008).

Thus, given the studies of latitudinal clines and adaptation from *Drosophila* and the nascent understanding of clock gene contributions to behavioral phenotypes in present day humans, the directional effects of introgressed circadian gene variants toward early chronotype and decreased gene regulation we observed can be viewed as potentially adaptive. More complex chronotype phenotyping and mechanistic studies of the variants of interest are needed to fully understand these observations.

Finally, to explore evidence for positive selection on introgressed variants in AMH, we analyzed results from three recent methods for detecting adaptive introgression. All methods identified circadian loci as candidates for adaptive introgression. However, we note that the predictions of these methods have only modest overlap with one another, underscoring the difficulty of identifying adaptive introgression. Nonetheless, many of these loci, especially those supported by both Racimo and *MaLAdapt*, are good candidates for adaptive introgression given their functional associations with circadian genes Several limitations must be considered when interpreting our results. First, it is challenging to quantify the complexity of traits with a large behavioral component (like chronotype) and infer their variation from genomic information alone. Nevertheless, we believe our approach of focusing on molecular aspects (splicing, gene regulation) of genomic loci with relevance to circadian biology, in parallel to GWAS-based associations, lends additional support to the divergence in chronotype between archaic hominins and modern humans. Second, we also note that circadian rhythms contribute to many biological systems, so the variants in these genes tend to be associated with a variety of phenotypes. Thus, there is also the potential that selection acted on other phenotypes influenced by circadian variation than those related directly to chronotype. Third, given the complexity of circadian biology, there is no gold standard set of circadian genes. We focus on the core clock genes and a broader set of expert-curated genes relevant to circadian systems, but it is certainly possible that other genes with circadian effects are not considered. Fourth, recent adaptive evolution is challenging to identify, and this is especially challenging for introgressed loci. Nonetheless, we find several circadian loci with evidence of adaptive introgression from more than one method. Finally, given the many environmental factors that differed between African and non-African environments, it is difficult to definitively determine whether selection on a particular locus was the result of variation in light levels vs. other related factors, such as temperature. Nonetheless, given the observed modern associations with chronotype for many of these variants, we believe it is a plausible target.

In conclusion, studying how humans evolved in the face of changing environmental pressures is necessary to understanding variation in present-day phenotypes and the potential tradeoffs that influence propensity to different diseases in modern environments (Benton *et al*., 2021). Here, we show that genomic regions involved in circadian biology exhibited substantial functional divergence between separate hominin populations. Furthermore, we show that introgressed variants contribute to variation in AMH circadian phenotypes today in ways that are consistent with an adaptive benefit.

## METHODS

### Circadian gene selection

Circadian biology is a complex system due to its high importance in the functioning of biological timing in diverse biological systems. For that reason, determining which genes are crucial for selection to environment response related to light exposure is not a straight forward process. To address this issue, we look at different sources of genome annotation databases and searched for genes and variants associated with circadian related phenotypes. We considered all human protein-coding genes in the Gene Ontology database annotated with the GO:0007623 (“circadian rhythm”) term or terms annotated with relationship “is_a”, “part_of”, “occurs_in”, or “regulates” circadian rhythm. We also considered genes containing experimental or orthologous evidence of circadian function in the Circadian Gene Database (CGDB), the GWAS Catalog genes containing “chronotype” or “circadian rhythm” associated variants, and a curated set of genes available in WikiPathways [https://www.wikipathways.org/index.php/Pathway:WP3594, https://doi.org/10.1093/nar/gkaa1024]. The final set of circadian genes was curated by Dr. Douglas McMahon.

To select the candidate circadian genes with the highest confidence, we defined a hierarchy system where genes annotated by McMahon or annotated in 3 out of 4 other sources receive a “High” level of confidence. Genes with evidence from 2 out of 4 of the sources are assigned a “Medium” level of confidence. Genes annotated as circadian only in 1 out of 4 sources are assigned to Low confidence and not considered in our circadian gene set. We then defined our set of circadian variants from the 1000 Genomes Project using the official list of circadian genes. The variants are included in analysis of coding, non-coding, regulatory, eQTL, human-specific, archaic-specific, and introgressed variants.

### Definition of lineage-specific variants

To identify candidate variants that are specific to the human and the archaic lineages, we used a set of variants published by Kuhlwilm and Boeckx (Kuhlwilm and Boeckx, 2019) (https://doi.org/10.1038/s41598-019-44877-x). The variants were extracted from the high-coverage genomes of three archaics: a 122,000-year-old Neanderthal from the Altai Mountains (52x coverage), a 52,000-year-old Neanderthal from Vindija in Croatia (30x coverage), and a 72,000-year-old Denisovan from the Altai Mountains (30x coverage). The variants were called in the context of the human genome hg19/GRCh37 reference. The total number of variant sites after applying filters for high coverage sites and genotype quality is 4,437,803. A human-specific variant is defined as a position where all the humans in the 1000 Genomes Project carry the derived allele and all the archaics carry the ancestral allele. An archaic-specific is defined as a position where all the archaics carry the derived allele and the derived allele is absent or extremely rare (<= 0.00001) across all human populations. Note that introgressed archaic alleles are not included in the “archaic-specific” set. These criteria resulted in 9,424 human specific and 33,184 archaic-specific variants.

### Enrichment of lineage-specific variants among functional regions of the genome

We intersected the sets of lineage-specific variants with several sets of annotated functional genomic regions. Inside circadian gene regions (Gencode v29), we found 156 human-specific variants and 341 archaic-specific variants. In circadian promoter regions, we found 6 human-specific variants and 19 archaic-specific variants. Promoters were defined as regions 5 kb up- to 1 kb downstream from a transcription start site. In distal regulatory elements, we found 247 human-specific variants and 807 archaic-specific variants. For this last set, we considered candidate cis-regulatory elements (cCREs) published by ENCODE (Moore *et al*., 2020) within 1 Mb of the circadian genes.

To compute whether lineage-specific variants are more abundant than expected in circadian genes, we applied a Fisher’s exact test to the sets of human- and archaic-specific variants in regulatory, promoter, and gene regions. Human and archaic-specific variants are significantly enriched in both regulatory (Human: OR=1.25, P=8.39e-4; Archaic: OR=1.16, P=6.15e-5) and gene (Human: OR=1.84, P=7.06e-12; Archaic: OR=1.13, P=0.023) regions. The enrichment observed in the promoters of both lineages is not supported by a significant p-value (Human: OR=1.21, P=0.65; Archaic: OR=1.09, P=0.63).

### Genes containing archaic variants with evidence of alternative splicing

We used a set of archaic variants annotated with the splice altering probabilities to identify circadian genes that may be differentially spliced between archaic hominins and AMH (Brand, Colbran and Capra, 2023). We considered variants from four archaic individuals: the Altai, Chagyrskaya, and Vindija Neanderthals and the Altai Denisovan. These archaic variants were annotated using SpliceAI (Jaganathan *et al*., 2019) and we considered any variant with a maximum delta, or splice altering probability, > 0.2. We identified 36 archaic-specific splice altering variants, defined as those variants absent from 1000 Genomes Project, among 28 circadian genes. Next, we tested for enrichment among this gene set using an empirical null approach (McArthur *et al*., 2022; Brand, Colbran and Capra, 2023). We shuffled the maximum deltas among 1,607,350 variants 10,000 times and counted the number of circadian genes with a splice altering variant each iteration. Enrichment was calculated as the number of observed genes (N = 28) divided by the mean gene count among 10,000 shuffles. In addition to all genes with archaic-specific variants, we considered six other subsets among these variants: 1) genes with variants private to the Altai Neanderthal, 2) genes with variants private to the Chagyrskaya Neanderthal, 3) genes with variants private to the Altai Denisovan, 4) genes with variants private to all Neanderthals, 5) genes with variants shared among all archaic individuals, and 6) genes with variants private to the Vindija Neanderthal. Finally, we considered a subset of splice altering variants that were identified as tag SNPs by Vernot et al. (Vernot *et al*., 2016).

### PrediXcan

To understand the difference in circadian biology between present-day humans and archaic hominins, we analyzed predictions on gene regulation. We considered the results from PrediXcan gene regulation predictions across 44 tissues from the PredictDB Data Repository (http://predictdb.org/). The models were trained on GTEx V6 using variants identified in 2,504 present-day humans in the 1000 Genomes Project phase 3 within 1 Mb of each circadian gene. The original analysis includes predictions for 17,748 genes for which the models explained a significant amount of variance in gene expression in each tissue (FDR < 0.05). The prediction models were also applied to the Altai and Vindija Neanderthals and the Denisovan. The resulting predictions are normalized values of the distribution observed in GTEx individuals used to train the original prediction models. Each prediction contains an empirical P-value which was calculated for each gene and tissue pair to define genes that are divergently regulated between archaic hominins and humans. The P-value is obtained by calculating the proportion of humans from the 1000 Genomes Project that have predictions more extreme compared to the human median than the archaic individual. Significantly DR genes are defined as those where the archaic prediction falls outside the distribution of humans in the 1000 Genomes Project predictions.

We tested whether the circadian genes in our set are more likely to be DR compared to an empirical null distribution from random gene sets of the same size. We account for the fact that some genes are modeled in more tissues than others by matching the distribution of tissues in which each gene could be modeled in the random sets to our set. Among 1,467 DR genes in the Altai Neanderthal we find 23 DR circadian genes out of the total 236 genes in the circadian set. We iterate through the permutation analysis 1,000,000 times and find an enrichment of 1.21 (P=0.19). A similar analysis is done in the Vindija Neanderthal (1,536 total DR, 21 circadian DR, enrichment of 1.05, P=0.43) and the Denisovan individual (1,214 total DR, 19 circadian DR, enrichment of 1.20, P=0.24). In this study, we define a set of DR genes as the intersection between DR genes in all three archaics, resulting in a set of 16 genes.

### Enrichment of introgressed variants in eQTL

We performed an enrichment analysis using Pearson’s chi-squared test to evaluate if there is overrepresentation of introgressed alleles in our set of circadian variants using the GTEx dataset. We did a liftOver of the GTEx v8 dataset from hg38 to hg19. The original hg38 set contains 4,631,659 eQTLs across 49 tissues. After the LiftOver, 4,608,446 eQTLs remained, with the rest not mapping. We used the archaic introgressed variants dataset from Browning 2018. The set contains 863,539 variants that are introgressed in humans originating in archaic hominins. We performed an intersection between the set of genes containing evidence for eQTLs and our set of 246 circadian genes to retrieve a subset of variant sites with evidence of being eQTL in circadian genes. The resulting subset contained 97,441 circadian eQTLs in 49 tissues and 239 genes. We further intersected the introgressed variants and the set of eQTL, resulting in 128,138 introgressed eQTLs. The final set of eQTLs that are circadian and also introgressed is 3,857.

### Direction of effect of chronotype associations

To explore the effect of archaic introgression in circadian dreams on human chronotype, we quantified the direction of effect of variants associated to a Morning/Evening person trait in a GWAS analysis of the UK Biobank (http://www.nealelab.is/uk-biobank/). The variants were LD clumped using PLINK v1.9 (R^2^>0.5). We generated cumulative proportion values on the beta values assigned to each associated variant on an ascending order of P-values.

### Detection of latitudinal clines in chronotype associations

To evaluate latitudinal clines in chronotype-associated variants, we assigned a latitude to each of the Eurasian 1000 Genomes Project populations. The latitude of diaspora populations was set to their ancestral country (GIH Gandhinagar in Gujarat: 23.223, STU Sri Jayawardenepura Kotte: 6.916667, ITU Amaravati in Andhra Pradesh: 16.5131, CEU: 52.372778). CEU was assigned a latitude in Amsterdam, following an analysis that shows that this group is more closely related to Dutch individuals (Lao *et al*., 2008). We then used the LDlink API to retrieve allele frequencies for each introgressed morningness variant in Eurasian individuals (Machiela and Chanock, 2015). Variants that follow a latitudinal cline were identified using linear regression statistics requiring correlation coefficient (R >= 0.65) and P-value (P <= 0.5).

### Detection of pleiotropy in the set of introgressed circadian variants

To understand the extent of different phenotypes associated with the introgressed circadian variants, we first extracted genome-wide associations from Open Targets Genetics (https://genetics.opentargets.org/) for each of the variants with evidence of introgression (Browning *et al*., 2018). Only the variants with significant p-values were analyzed. The p-value threshold was set at the genome-wide significance level (P=5e-8). We split the variants in two sets: introgressed circadian and introgressed non-circadian. Many of these variants are not associated with any phenotype. We performed a Fisher’s exact test to analyze which of the two sets contains a higher ratio of SNPs with at least one association versus SNPs with no association. The result showed that the circadian set had a significantly higher ratio (OR=1.36, P=5e-29). Then we calculated the total of unique traits associated with each of the variants, given that the SNP has at least one association. We used a Mann-Whitney U test to understand which set is represented by a higher level of traits per SNP. The circadian set was slightly more pleiotropic, and the result is supported by a significant p-value (P=5.4e-3).

### Identifying introgressed circadian variants with evidence of adaptive introgression

We sought to identify circadian variants that contain evidence of adaptive introgression (AI). To achieve this, we collected AI predictions from a method that applied various summary statistics on 1000 Genomes Project data (Racimo, Marnetto and Huerta-Sánchez, 2017) and two sets of genomic regions that were measured for their likelihood to be under AI by two machine learning methods: *genomatnn* and *MaLAdapt*. *genomatnn* is a convolutional neural network trained to identify adaptive introgression based on simulations (Gower *et al*., 2021). *MaLAdapt* is a machine learning algorithm trained to find adaptive introgression based on simulations using an extra-trees classifier (ETC) (Zhang *et al*., 2023). Following the thresholds used in each paper, a region is considered to be under AI if the prediction value assigned to it meets a threshold of 0.5 or 0.9, respectively. To find the variants of interest that fall into AI regions, we intersected the set of introgressed circadian SNPs with the Racimo et al. 2015, *genomatnn* and the *MaLAdapt* regions individually. The set of introgressed circadian variants contains variants inside circadian genes, in circadian promoter regions (5 kb up- and 1 kb downstream of the TSS), and variants with regulatory function (cCREs) flanking circadian genes by 1 Mb.

## DATA AVAILABILITY

The data underlying this article are available in the article and in its online supplementary material.

## ACKNOWLEDGMENTS

We thank members of the Capra Lab for helpful comments on this work. This work was conducted in part using the resources of the Advanced Computing Center for Research and Education at Vanderbilt University, Nashville, TN. This work was supported by the National Institutes of Health [R35GM127087 to JAC, R01GM117650 to DM, F30HG011200 to EM, T32GM080178 to Vanderbilt University (EM), and T32HG009495 to the University of Pennsylvania (LLC)].

## AUTHOR CONTRIBUTIONS

Conceptualization: KV, JAC; Methodology: KV, LC, EM, CB, DR, JS, DM, JAC; Investigation: KV, LC, EM, CB, JAC; Writing – Original Draft: KV, JAC; Writing – Review & Editing: KV, LC, EM, CB, DR, DM, JAC; Funding Acquisition: JAC; Resources: JAC; Supervision: JAC.

