## Supplemental figures and tables for "Archaic Introgression Shaped Human Circadian Traits"

**SUPPLEMENTARY MATERIAL**

**SUPPLEMENTARY FIGURES**

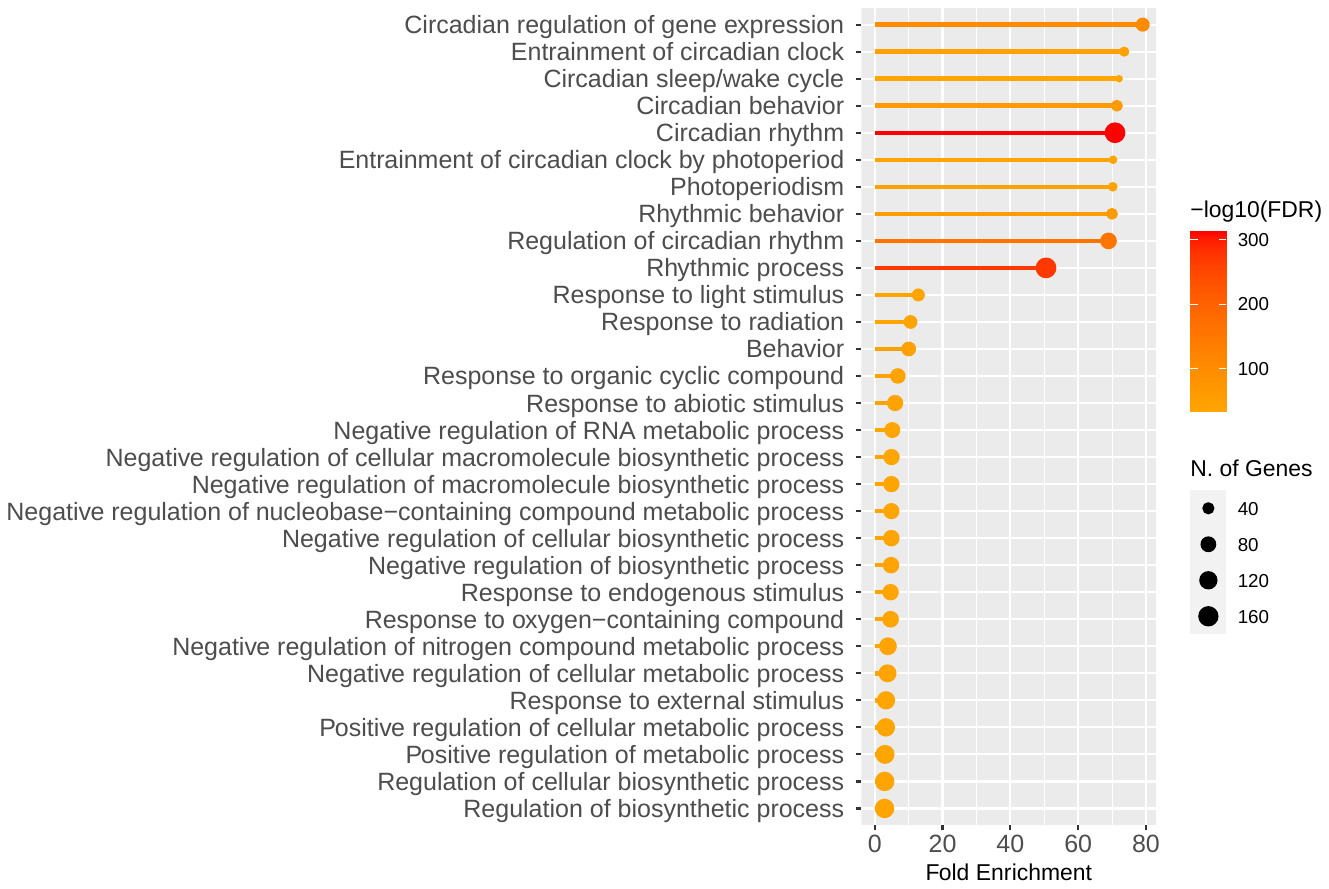

**Figure S1**. **GO terms associated with the 246 circadian genes.** Generated by ShinyGO, <http://bioinformatics.sdstate.edu/go75/> . The strong enrichment for circadian terms supports their relevance, but we note that GO annotations were used to select some of these genes, so this should not be viewed as independent.

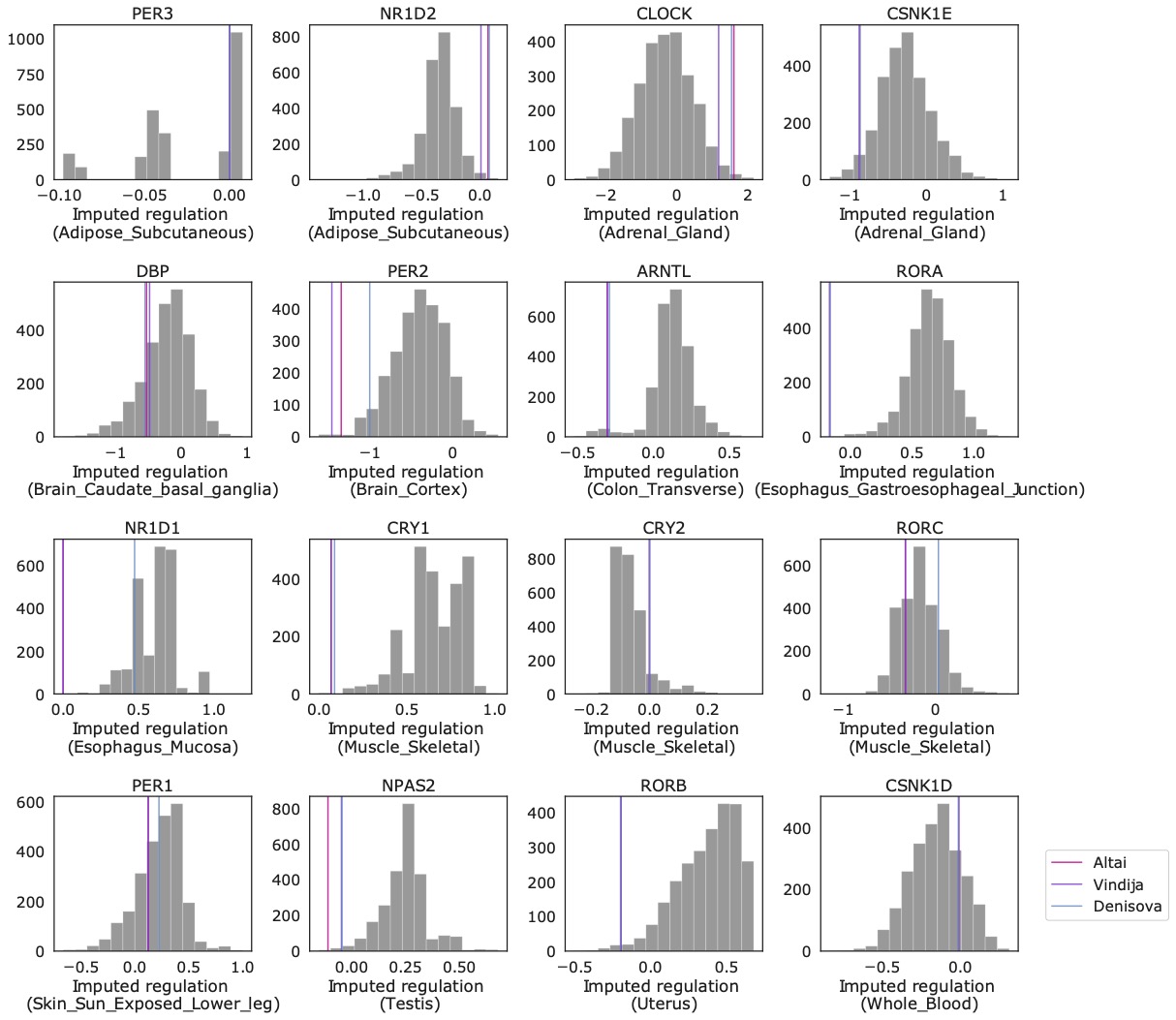

**Figure S2.** **Comparison of imputed regulation for** **core circadian genes between modern humans and archaic hominins.** Comparison of the imputed regulation of core circadian genes between 2504 humans in 1000 Genomes Phase 3 (gray bars) and three archaic individuals (vertical lines). For each core circadian gene, the tissue with the lowest average P-value for archaic difference from humans is plotted. Archaic gene regulation is at the extremes of the human distribution for several core genes: *CRY1*, *PER2*, *NPAS2,* *NR1D2*, *RORA*.

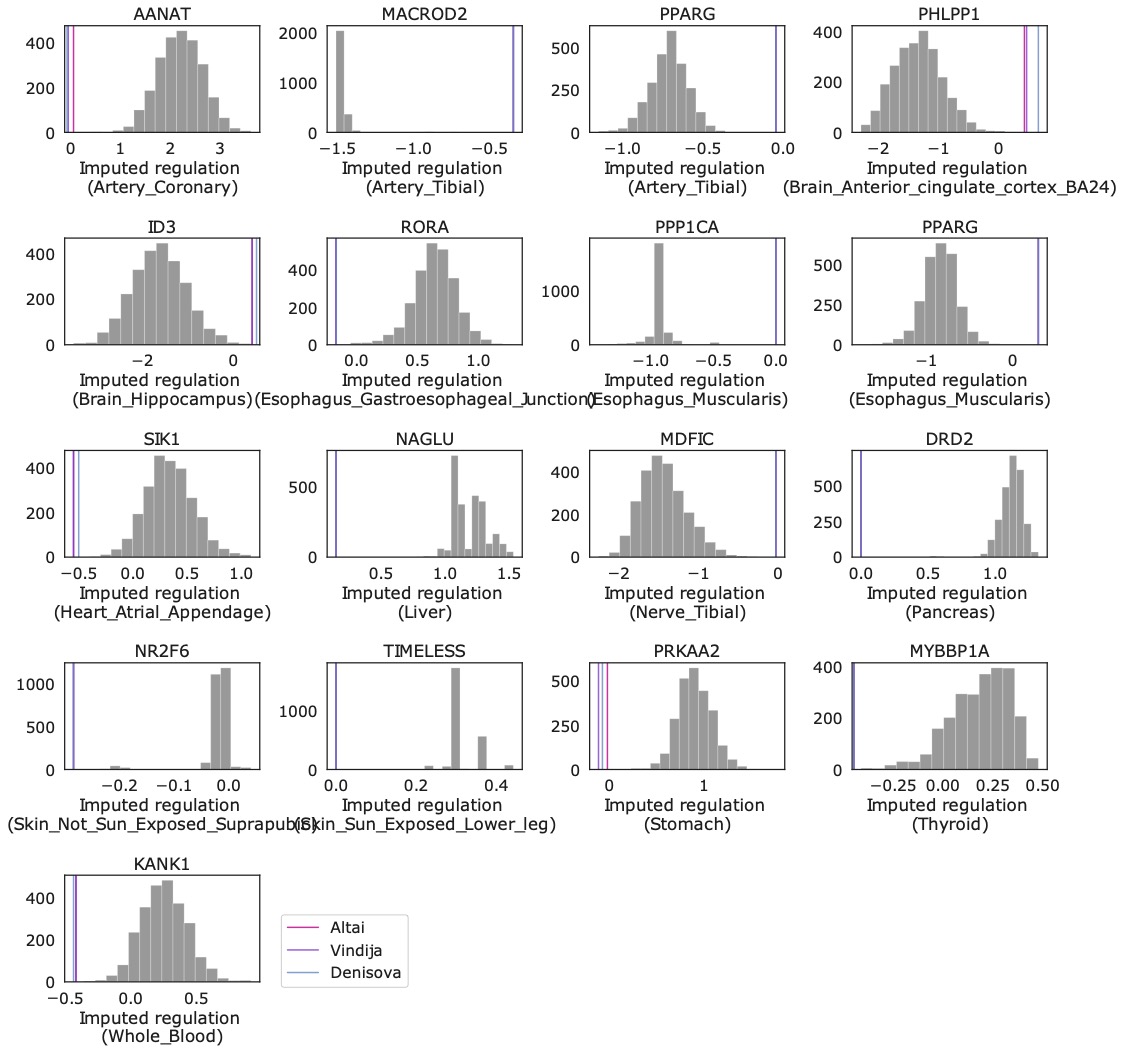

**Figure S3**. **Distributions of gene regulation predictions in all divergently regulated circadian genes.** Divergent regulation indicates that the archaic individuals (colored lines) each had imputed regulation more extreme than all 2504 modern humans from the 1000 Genomes Project (gray bars). For each gene, the tissue with largest average archaic difference from humans is plotted.

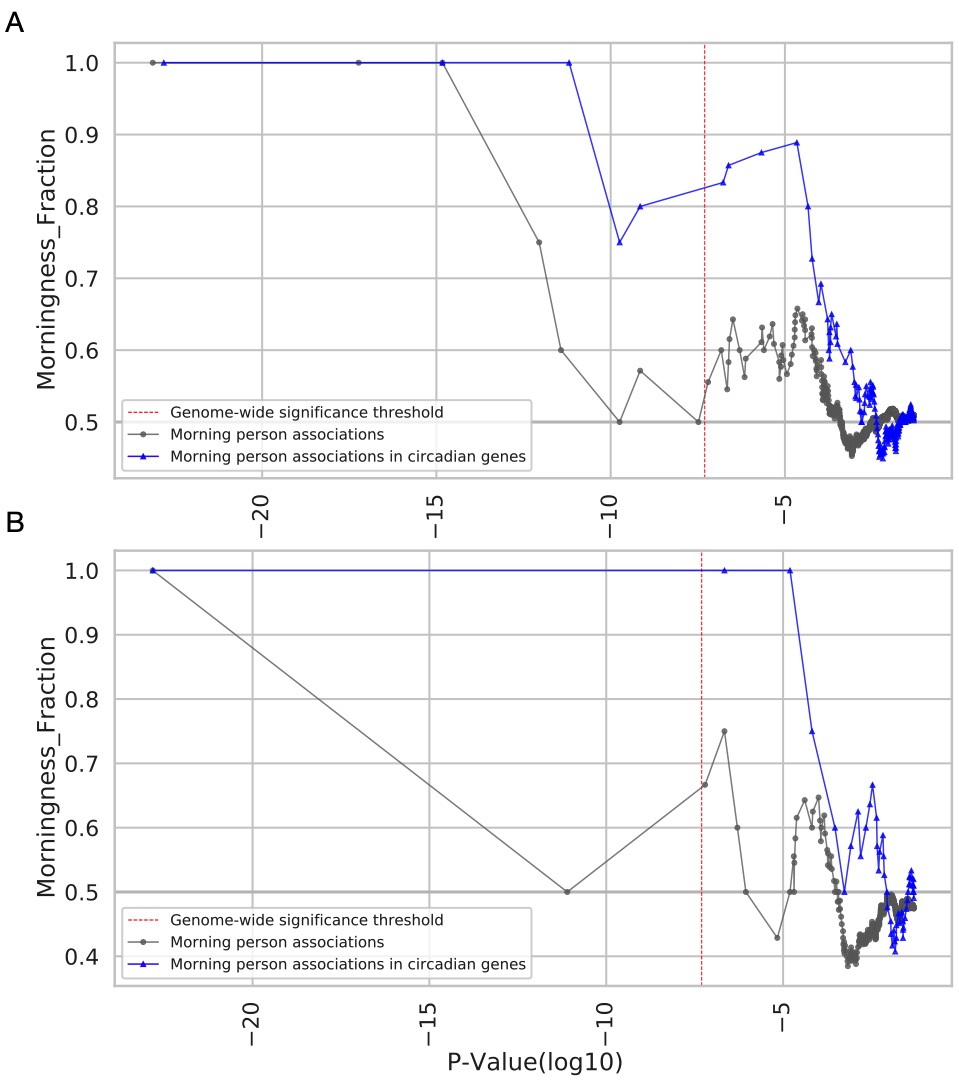

**Figure S4. Introgressed variants associate with increased morningness.** A) Cumulative fraction of morningness increasing variants reported as introgressed by Browning et al. 2018, and at least one other introgression detection method. B) Cumulative fraction of morningness increasing variants reported as introgressed by Browning et al. 2018 and five other introgression detection methods.

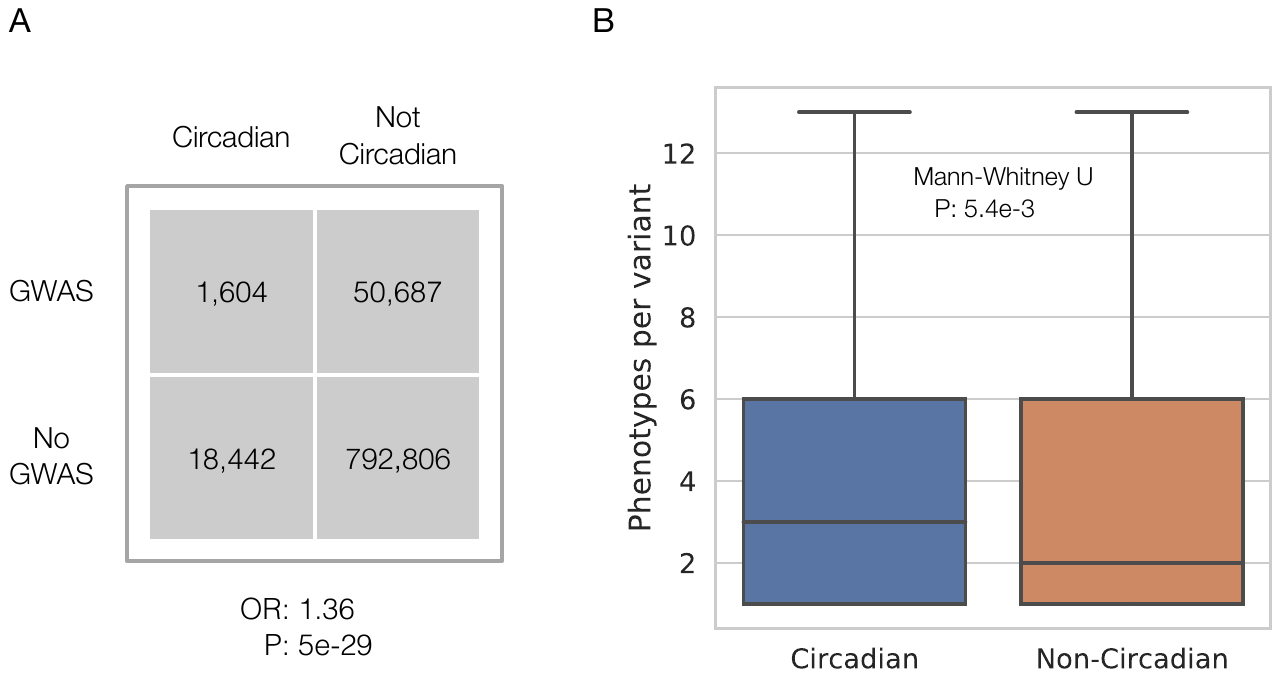

**Figure S5. Pleiotropy in introgressed circadian variants.** **A)** The relationship between GWAS associations and circadian introgressed variants versus non-circadian introgressed variants. The circadian set contains significantly more sites with at least one association than the non-circadian set (Fisher’s exact: OR=1.36, P=5e-29). The GWAS associations were retrieved from Open Targets Genetics and were filtered by the genome-wise significance (P=5e-8). **B)** The distribution of phenotype associations per variant in the set of circadian and non-circadian introgressed variants. The circadian set shows significantly higher pleiotropy (Mann-Whitney U test: P=5.4e-3). Outliers (beyond 1.5 times the interquartile range) were removed for visualization; see Table S9.

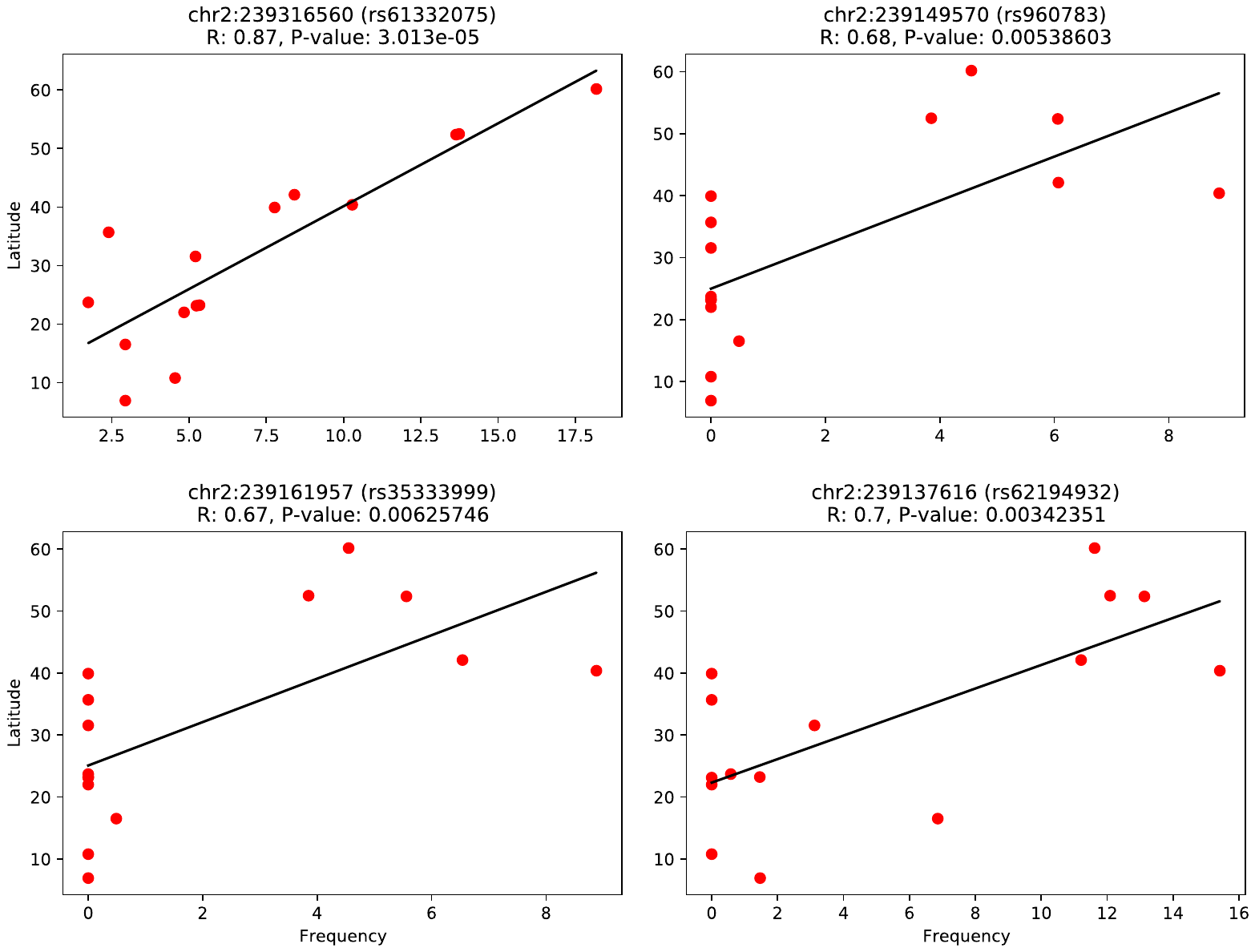

**Figure S6. Introgressed haplotypes associated with morningness follow a latitudinal cline**. rs61332075 is the leading SNP in a large chromosome 2 haplotype that contains a SNP (rs75804782) previously reported to show a latitudinal cline in Eurasia. rs960783 and rs35333999 are tag SNPs in a nearby haplotype that also shows a latitudinal cline in Eurasia. rs62194932 is in moderate LD (R^2^ of ~0.35 in EUR) with this haplotype and shows a similar latitudinal cline. Red dots represent 1000 Genomes Project populations of Eurasian ancestry.

**SUPPLEMENTARY TABLES**

**Table S1**. List of circadian genes analyzed in this study and the sources of evidence reporting the gene's relevance in circadian cycles. See Supplementary Tables Excel file.

**Table S2**. Lineage specific variants inside circadian genes, in circadian promoter regions, and with regulatory function flanking circadian genes by 1Mb. See Supplementary Tables Excel file.

**(cont’d)**

**Table S3**. Genes containing variants predicted to be splice-altering by the SpliceAI method in the Altai Neanderthal (A), Vindija Neanderthal (V), Chagyrskaya Neanderthal (C), and Denisova (D).

| **GeneID** | **GeneName** | **Description** | **Locus** | **A** | **C** | **D** | **V** |
| --- | --- | --- | --- | --- | --- | --- | --- |
| ENSG00000153064 | BANK1 | B cell scaffold protein with ankyrin repeats 1 | chr4_102911885 | 0 | 0 | 1 | 0 |
| ENSG00000158941 | CCAR2 | cell cycle and apoptosis regulator 2 | chr8_22472265 | 0 | 0 | 0 | 1 |
| ENSG00000107736 | CDH23 | cadherin related 23 | chr10_73298326 | 1 | 1 | 1 | 1 |
|  |  |  | chr10_73405409 | 1 | 1 | 0 | 1 |
|  |  |  | chr10_73466969 | 0 | 1 | 0 | 0 |
|  |  |  | chr10_73487652 | 0 | 0 | 1 | 0 |
| ENSG00000134852 | CLOCK | clock circadian regulator | chr4_56296172 | 0 | 0 | 1 | 0 |
| ENSG00000105662 | CRTC1 | CREB regulated transcription coactivator 1 | chr19_18867740 | 1 | 1 | 0 | 1 |
| ENSG00000057593 | F7 | coagulation factor VII | chr13_113769975 | 0 | 0 | 1 | 0 |
| ENSG00000161040 | FBXL13 | F-box and leucine rich repeat protein 13 | chr7_102667940 | 1 | 1 | 0 | 1 |
| ENSG00000119771 | KLHL29 | kelch like family member 29 | chr2_23805847 | 1 | 1 | 0 | 1 |
| ENSG00000205213 | LGR4 | leucine rich repeat containing G protein-coupled receptor 4 | chr11_27443527 | 0 | 0 | 1 | 0 |
| ENSG00000005810 | MYCBP2 | MYC binding protein 2 | chr13_77853722 | 0 | 0 | 1 | 0 |
| ENSG00000140396 | NCOA2 | nuclear receptor coactivator 2 | chr8_71180317 | 0 | 0 | 1 | 0 |
| ENSG00000141027 | NCOR1 | nuclear receptor corepressor 1 | chr17_16040743 | 0 | 0 | 1 | 0 |
| ENSG00000064300 | NGFR | nerve growth factor receptor | chr17_47581270 | 0 | 0 | 1 | 0 |
| ENSG00000204640 | NMS | neuromedin S | chr2_101097107 | 1 | 0 | 0 | 0 |
| ENSG00000132326 | PER2 | period circadian regulator 2 | chr2_239187089 | 0 | 0 | 0 | 1 |
| ENSG00000149177 | PTPRJ | protein tyrosine phosphatase receptor type J | chr11_48009909 | 0 | 0 | 0 | 1 |
|  |  |  | chr11_48041749 | 0 | 0 | 1 | 0 |
| ENSG00000173482 | PTPRM | protein tyrosine phosphatase receptor type M | chr18_7666317 | 0 | 0 | 0 | 1 |
|  |  |  | chr18_7666418 | 0 | 1 | 0 | 0 |
| ENSG00000152061 | RABGAP1L | RAB GTPase activating protein 1 like | chr1_174606762 | 0 | 1 | 0 | 0 |
|  |  |  | chr1_174876469 | 1 | 0 | 0 | 0 |
| ENSG00000173933,  ENSG00000173914 | RBM4,  RBM4B | RNA binding motif protein 4,  RNA binding motif protein 4B | chr11_66433175 | 1 | 1 | 0 | 1 |
| ENSG00000141576 | RNF157 | ring finger protein 157 | chr17_74152940 | 1 | 1 | 0 | 1 |
| ENSG00000134318 | ROCK2 | Rho associated coiled-coil containing protein kinase 2 | chr2_11370990 | 0 | 0 | 1 | 0 |
|  |  |  | chr2_11484478 | 0 | 0 | 0 | 1 |
| ENSG00000198963 | RORB | RAR related orphan receptor B | chr9_77113671 | 0 | 0 | 0 | 1 |
| ENSG00000143365 | RORC | RAR related orphan receptor C | chr1_151796494 | 1 | 1 | 0 | 1 |
| ENSG00000103546 | SLC6A2 | solute carrier family 6 member 2 | chr16_55707020 | 0 | 0 | 1 | 0 |
| ENSG00000108576 | SLC6A4 | solute carrier family 6 member 4 | chr17_28547727 | 1 | 1 | 0 | 1 |
| ENSG00000072310 | SREBF1 | sterol regulatory element binding transcription factor 1 | chr17_17719515 | 1 | 0 | 0 | 0 |
|  |  |  | chr17_17719517 | 1 | 0 | 0 | 0 |
| ENSG00000141510 | TP53 | tumor protein p53 | chr17_7578263 | 0 | 0 | 1 | 0 |
| ENSG00000140836 | ZFHX3 | zinc finger homeobox 3 | chr16_72966622 | 0 | 1 | 0 | 1 |

**Table S4**. Genes predicted to be divergently regulated (DR) in the archaics from Altai (A), Vindija (V), and Denisova (D) by the PrediXcan method.

| **GeneID** | **GeneName** | **Description** | **GTEx Tissue** | **A** | **V** | **D** |
| --- | --- | --- | --- | --- | --- | --- |
| ENSG00000129673 | AANAT | aralkylamine N-acetyltransferase | Artery_Coronary | 1 | 1 | 1 |
| ENSG00000174080 | CTSF | cathepsin F | Adipose_Visceral_Omentum | 1 | 0 | 1 |
|  |  |  | Brain_Cerebellar_Hemisphere | 1 | 0 | 1 |
| ENSG00000149295 | DRD2 | dopamine receptor D2 | Pancreas | 1 | 1 | 1 |
| ENSG00000107485 | GATA3 | GATA binding protein 3 | Skin_Sun_Exposed_Lower_leg | 0 | 1 | 0 |
| ENSG00000115738 | ID2 | inhibitor of DNA binding 2 | Skin_Not_Sun_Exposed_Suprapubic | 1 | 0 | 0 |
| ENSG00000117318 | ID3 | inhibitor of DNA binding 3, HLH protein | Brain_Hippocampus | 1 | 1 | 1 |
| ENSG00000143772 | ITPKB | inositol-trisphosphate 3-kinase B | Esophagus_Mucosa | 1 | 0 | 0 |
| ENSG00000107104 | KANK1 | KN motif and ankyrin repeat domains 1 | Whole_Blood | 1 | 1 | 1 |
| ENSG00000172264 | MACROD2 | mono-ADP ribosylhydrolase 2 | Artery_Tibial | 1 | 1 | 1 |
| ENSG00000135272 | MDFIC | MyoD family inhibitor domain containing | Nerve_Tibial | 1 | 1 | 1 |
| ENSG00000132382 | MYBBP1A | MYB binding protein 1a | Muscle_Skeletal | 1 | 0 | 1 |
|  |  |  | Thyroid | 1 | 1 | 1 |
| ENSG00000108784 | NAGLU | N-acetyl-alpha-glucosaminidase | Liver | 1 | 1 | 1 |
| ENSG00000126368 | NR1D1 | nuclear receptor subfamily 1 group D member 1 | Adipose_Subcutaneous | 0 | 1 | 0 |
|  |  |  | Esophagus_Mucosa | 1 | 0 | 1 |
| ENSG00000160113 | NR2F6 | nuclear receptor subfamily 2 group F member 6 | Skin_Not_Sun_Exposed_Suprapubic | 1 | 1 | 1 |
| ENSG00000140538 | NTRK3 | neurotrophic receptor tyrosine kinase 3 | Adipose_Visceral_Omentum | 1 | 0 | 1 |
| ENSG00000081913 | PHLPP1 | PH domain and leucine rich repeat protein phosphatase 1 | Brain_Anterior_cingulate_cortex_BA24 | 1 | 1 | 1 |
| ENSG00000132170 | PPARG | peroxisome proliferator activated receptor gamma | Esophagus_Muscularis | 1 | 1 | 1 |
|  |  |  | Artery_Tibial | 1 | 1 | 1 |
| ENSG00000172531 | PPP1CA | protein phosphatase 1 catalytic subunit alpha | Esophagus_Muscularis | 1 | 1 | 1 |
| ENSG00000162409 | PRKAA2 | protein kinase AMP-activated catalytic subunit alpha 2 | Stomach | 1 | 1 | 1 |
| ENSG00000069667 | RORA | RAR related orphan receptor A | Esophagus_Gastroesophageal_Junction | 1 | 1 | 1 |
| ENSG00000142178 | SIK1 | salt inducible kinase 1 | Heart_Atrial_Appendage | 1 | 1 | 1 |
| ENSG00000111602 | TIMELESS | timeless circadian regulator | Skin_Sun_Exposed_Lower_leg | 1 | 1 | 1 |
| ENSG00000141510 | TP53 | tumor protein p53 | Brain_Nucleus_accumbens_basal_ganglia | 1 | 0 | 1 |
| ENSG00000140836 | ZFHX3 | zinc finger homeobox 3 | Cells_Cultured_fibroblasts | 1 | 0 | 1 |
|  |  |  | Small_Intestine_Terminal_Ileum | 0 | 1 | 1 |

**Table S5**. Introgressed variants (Browning et al. 2018) with evidence of being eQTL (GTEx) inside circadian genes, circadian promoter region, or with regulatory function flanking circadian genes by 1Mb. See Supplementary Tables Excel file.

**(cont’d)**

**Table S6.** Counts of circadian genes expressed in each GTEx tissue (TPM >= 1).

| **Tissue** | **Gene_count** | **Percentage** |
| --- | --- | --- |
| Testis | 206 | 83.74 |
| Pituitary | 197 | 80.08 |
| Brain_Nucleus_accumbens_basal_ganglia | 195 | 79.27 |
| Brain_Hypothalamus | 195 | 79.27 |
| Brain_Frontal_Cortex_BA9 | 195 | 79.27 |
| Brain_Cortex | 195 | 79.27 |
| Brain_Caudate_basal_ganglia | 193 | 78.46 |
| Brain_Anterior_cingulate_cortex_BA24 | 191 | 77.64 |
| Lung | 190 | 77.24 |
| Brain_Amygdala | 189 | 76.83 |
| Brain_Putamen_basal_ganglia | 188 | 76.42 |
| Fallopian_Tube | 187 | 76.02 |
| Brain_Hippocampus | 187 | 76.02 |
| Prostate | 187 | 76.02 |
| Cervix_Endocervix | 186 | 75.61 |
| Vagina | 186 | 75.61 |
| Breast_Mammary_Tissue | 185 | 75.2 |
| Brain_Cerebellum | 185 | 75.2 |
| Cervix_Ectocervix | 184 | 74.8 |
| Small_Intestine_Terminal_Ileum | 184 | 74.8 |
| Brain_Substantia_nigra | 184 | 74.8 |
| Nerve_Tibial | 183 | 74.39 |
| Thyroid | 183 | 74.39 |
| Adipose_Subcutaneous | 181 | 73.58 |
| Artery_Coronary | 180 | 73.17 |
| Brain_Cerebellar_Hemisphere | 179 | 72.76 |
| Colon_Transverse | 179 | 72.76 |
| Bladder | 179 | 72.76 |
| Skin_Sun_Exposed_Lower_leg | 178 | 72.36 |
| Adipose_Visceral_Omentum | 177 | 71.95 |
| Ovary | 176 | 71.54 |
| Spleen | 175 | 71.14 |
| Artery_Tibial | 175 | 71.14 |
| Skin_Not_Sun_Exposed_Suprapubic | 175 | 71.14 |
| Uterus | 175 | 71.14 |
| Brain_Spinal_cord_cervical_c-1 | 175 | 71.14 |
| Stomach | 173 | 70.33 |
| Esophagus_Gastroesophageal_Junction | 172 | 69.92 |
| Kidney_Medulla | 172 | 69.92 |
| Colon_Sigmoid | 172 | 69.92 |
| Adrenal_Gland | 172 | 69.92 |
| Minor_Salivary_Gland | 171 | 69.51 |
| Esophagus_Muscularis | 171 | 69.51 |
| Kidney_Cortex | 170 | 69.11 |
| Heart_Atrial_Appendage | 170 | 69.11 |
| Artery_Aorta | 169 | 68.7 |
| Esophagus_Mucosa | 165 | 67.07 |
| Pancreas | 164 | 66.67 |
| Liver | 162 | 65.85 |
| Heart_Left_Ventricle | 158 | 64.23 |
| Muscle_Skeletal | 156 | 63.41 |
| Cells_Cultured_fibroblasts | 156 | 63.41 |
| Cells_EBV-transformed_lymphocytes | 150 | 60.98 |
| Whole_Blood | 141 | 57.32 |

**Table S7**. Enrichment analysis on the introgressed circadian variants with evidence of being eQTL in each GTEx tissue (Fisher’s exact test).

| **Tissue** | **OR** | **P-value** | **Variants** | **log10(OR)** | **Bonferroni** |
| --- | --- | --- | --- | --- | --- |
| Adipose_Subcutaneous | 1.66152116 | 2.71E-52 | 1735 | 0.220505877 | 0.00102 |
| Adipose_Visceral_Omentum | 1.476860866 | 9.13E-28 | 1269 | 0.169339583 | 0.00102 |
| Adrenal_Gland | 1.702549549 | 1.63E-33 | 736 | 0.23109976 | 0.00102 |
| Artery_Aorta | 1.416931434 | 1.48E-21 | 1164 | 0.151348835 | 0.00102 |
| Artery_Coronary | 1.173064989 | 1.72E-03 | 469 | 0.069322073 | <=0.05 |
| Artery_Tibial | 2.137431738 | 4.40E-117 | 1984 | 0.329892254 | 0.00102 |
| Brain_Amygdala | 1.563173117 | 9.38E-13 | 326 | 0.194007078 | 0.00102 |
| Brain_Anterior_cingulate_cortex_BA24 | 1.367795929 | 4.51E-08 | 377 | 0.136021307 | 0.00102 |
| Brain_Caudate_basal_ganglia | 1.936875177 | 1.14E-52 | 798 | 0.287101633 | 0.00102 |
| Brain_Cerebellar_Hemisphere | 1.13396327 | 5.97E-03 | 603 | 0.054598988 | <=0.05 |
| Brain_Cerebellum | 1.24382714 | 8.53E-08 | 827 | 0.094760029 | 0.00102 |
| Brain_Cortex | 1.787971179 | 8.41E-46 | 901 | 0.252360514 | 0.00102 |
| Brain_Frontal_Cortex_BA9 | 1.465715751 | 6.62E-15 | 550 | 0.166049755 | 0.00102 |
| Brain_Hippocampus | 1.497305427 | 9.35E-14 | 441 | 0.175310399 | 0.00102 |
| Brain_Hypothalamus | 1.077370207 | 2.06E-01 | 331 | 0.032364961 | >0.05 |
| Brain_Nucleus_accumbens_basal_ganglia | 1.048180951 | 3.56E-01 | 462 | 0.020436263 | >0.05 |
| Brain_Putamen_basal_ganglia | 1.032429183 | 5.37E-01 | 410 | 0.013860272 | >0.05 |
| Brain_Spinal_cord_cervical_c-1 | 2.608684479 | 1.44E-75 | 563 | 0.416421554 | 0.00102 |
| Brain_Substantia_nigra | 0.436786642 | 1.14E-17 | 86 | -0.359730652 | 0.00102 |
| Breast_Mammary_Tissue | 1.528807665 | 3.16E-29 | 1069 | 0.184352852 | 0.00102 |
| Cells_Cultured_fibroblasts | 1.647755942 | 1.91E-51 | 1820 | 0.216892886 | 0.00102 |
| Cells_EBV-transformed_lymphocytes | 0.531777911 | 6.51E-17 | 155 | -0.274269707 | 0.00102 |
| Colon_Sigmoid | 1.330811081 | 3.46E-12 | 825 | 0.124116408 | 0.00102 |
| Colon_Transverse | 1.577020134 | 1.01E-33 | 1093 | 0.197837238 | 0.00102 |
| Esophagus_Gastroesophageal_Junction | 1.54633457 | 3.40E-31 | 1100 | 0.189303465 | 0.00102 |
| Esophagus_Mucosa | 1.010777799 | 7.61E-01 | 1146 | 0.004655695 | >0.05 |
| Esophagus_Muscularis | 2.043355917 | 1.92E-100 | 1751 | 0.31034402 | 0.00102 |
| Heart_Atrial_Appendage | 2.231494768 | 5.88E-110 | 1361 | 0.348595873 | 0.00102 |
| Heart_Left_Ventricle | 1.593604706 | 3.49E-32 | 961 | 0.202380604 | 0.00102 |
| Liver | 1.350488287 | 1.54E-08 | 447 | 0.130490822 | 0.00102 |
| Lung | 2.335072493 | 2.13E-139 | 1765 | 0.368300368 | 0.00102 |
| Minor_Salivary_Gland | 1.571257565 | 4.34E-14 | 358 | 0.196247382 | 0.00102 |
| Muscle_Skeletal | 1.255375135 | 3.87E-11 | 1403 | 0.098773522 | 0.00102 |
| Nerve_Tibial | 0.90900835 | 5.29E-03 | 1357 | -0.041432127 | <=0.05 |
| Ovary | 2.225014847 | 2.48E-55 | 565 | 0.347332913 | 0.00102 |
| Pancreas | 1.433186205 | 4.62E-20 | 951 | 0.156302619 | 0.00102 |
| Pituitary | 0.879911135 | 5.73E-03 | 557 | -0.055561187 | <=0.05 |
| Prostate | 1.365834247 | 3.60E-11 | 592 | 0.135397998 | 0.00102 |
| Skin_Not_Sun_Exposed_Suprapubic | 1.326473329 | 2.59E-16 | 1405 | 0.122698522 | 0.00102 |
| Skin_Sun_Exposed_Lower_leg | 1.387255205 | 1.29E-22 | 1642 | 0.142156363 | 0.00102 |
| Small_Intestine_Terminal_Ileum | 1.165873368 | 4.21E-03 | 413 | 0.066651382 | <=0.05 |
| Spleen | 0.685638014 | 4.39E-15 | 463 | -0.163905111 | 0.00102 |
| Stomach | 1.289923197 | 3.51E-09 | 719 | 0.110563853 | 0.00102 |
| Testis | 1.772493272 | 5.46E-66 | 1755 | 0.248584595 | 0.00102 |
| Thyroid | 1.683584451 | 9.27E-57 | 2010 | 0.226234906 | 0.00102 |
| Uterus | 1.145211042 | 5.54E-02 | 227 | 0.058885527 | >0.05 |
| Vagina | 1.106729697 | 1.53E-01 | 223 | 0.044041564 | >0.05 |
| Whole_Blood | 1.603439669 | 1.03E-43 | 1546 | 0.205052624 | 0.00102 |

**Table S8.** Introgressed non-circadian variants (Browning et al. 2018) associated with the Morning/evening person UK Biobank phenotype. See Supplementary Tables Excel file.

**Table S9.** Introgressed circadian variants (Browning et al. 2018) associated with the Morning/evening person UK Biobank phenotype. See Supplementary Tables Excel file.

**Table S10.** Trait associations for introgressed circadian variants from the Open Targets Genetics (https://genetics.opentargets.org) database, which combines GWAS data from the GWAS Catalog, UK Biobank, and several other sources. See Supplementary Tables Excel file.

**Table S11.** Introgressed SNPs (Browning et al. 2018) associated with at least 1 phenotype in the Open Targets database. See Supplementary Tables Excel file.

**Table S12.** Circadian variants located in regions predicted to be under adaptive introgression by genomatnn, MaLAdapt, or Racimo et al., 2017. See Supplementary Tables Excel file.

**Table S13.** Variants associated with morningness in the UKBiobank on chromosome 2 show a latitudinal cline in Eurasian populations. See Supplementary Tables Excel file.
